# *chrna3* modulates alcohol response

**DOI:** 10.1101/2024.08.22.607435

**Authors:** Joshua Raine, Caroline Kibat, Tirtha Das Banerjee, Antónia Monteiro, Ajay S. Mathuru

**Author notes:** At the time the work was performed (Caroline Kibat).

## Abstract

Alcohol use disorders (AUDs) are complex phenomena governed by genetics, neurophysiology, environment, and societal structures. New methods to understand the underlying neurogenetics are valuable for designing personalised interventional strategies. Here, we used a two-choice self-administration zebrafish assay (SAZA) to isolate the function of nicotinic acetylcholine receptor (nAChR) subunit alpha3 (*chrna3*) in alcohol response. Juvenile zebrafish, prior to sex differentiation, were examined in this study. They exhibited a biphasic response when self-administering alcohol that transitioned from attraction to aversion within minutes, suggesting they can regulate exposure to alcohol. This inverted U-shaped self-administration mirrored the effect alcohol has on shoaling behaviour. Exposure to low concentration of alcohol reduced anxiety-like behaviours, while sedative effects became prominent at higher concentrations resulting in reduced locomotion and uncoordinated swimming. In contrast, these responses are blunted in *chrna3* mutants. They exhibited prolonged alcohol self-administration, and increased gregariousness. Transcriptomic analyses suggest that glutamatergic and GABAergic neurotransmission alongside cholinergic signalling is impacted in the mutant brains. Our results thus suggest that *chrna3* dysfunction has a systemic change with an increase in alcohol tolerance being one effect. These findings also highlight the use of non-rodent alternatives to understand the neurogenetics of development of AUD.

## Introduction

Substance use disorders (SUDs), encompassing nicotine, alcohol, and opioid abuse, represent a significant global health burden, accounting for approximately 162.5 million disability-adjusted life years lost worldwide (Rehm and Shield, 2019a; Moeller and Terplan, 2020). According to the WHO, harmful alcohol use alone is estimated to have resulted in 5.3% of deaths across the world in 2022 (Rehm and Shield, 2019b; Witkiewitz et al., 2019; Aslam and Kwo, 2023). Efforts to understand the interplay among social, environmental, and biological factors to effectively counter alcohol use disorders (AUDs) continue to be a focus area, as the response to alcohol in vertebrates itself is a complex multifactorial phenomenon. Alcohol interacts with a range of neuromodulatory systems, engages reward processing circuits, and affects multiple neurotransmitter systems, including glutamate and GABA neurotransmission (Chvilicek et al., 2020). Both genetic and neurobiological factors contribute significantly to AUD development (Egervari et al., 2021; Koob and Vendruscolo, 2023). Thus, a comprehensive understanding of these factors, while accounting for individual variability, is crucial for developing effective AUD prevention and interventional strategies (Witkiewitz et al., 2019; Yang et al., 2022; Aslam and Kwo, 2023).

Animal models are indispensable for dissociating genetic contributors to the development of substance dependence in humans (Smith, 2020). While such studies are typically conducted in mammalian models, high operating costs can hinder studies on genetic variability and small, single gene effects (Cox, 2015; de Magalhães, 2015). Zebrafish present a cost-effective alternative, combining high throughput genetic manipulation with whole-brain live neural activity imaging (Cully, 2019). Additionally, anatomical and functional conservation with mammalian sub-cortical circuitry is now well established (Kily et al., 2008; Gerlai, 2020).

Zebrafish exhibit behavioural and physiological responses to psychoactive substances that are consistent with mammals, providing construct validity, however, studies have primarily employed indirect measures, like place preference or avoidance assays (Echevarria et al., 2011; Tsang et al., 2019). These are useful, but are considered inadequate to model SUD (Smith, 2020). Despite the difficulty in designing contingent, self-administration assays for aquatic animals, they are essential to gain predictive validity (Smith, 2020). Few studies have explored active choice (Schneider et al., 2023) or self-administration (Sterling et al., 2015; Bossé and Peterson, 2017; Nathan et al., 2022). Among these, studies with juveniles (∼4-5 weeks), as reported here, offer an advantage over both larvae or adults as behavioural responses become more consistent by this age, and a relatively large number of individuals can be examined to detect small effects.

Here, we expanded on the Self-Administration Zebrafish Assay (SAZA) (Nathan et al., 2022) to study *chrna3*’s role in acute, dynamic alcohol responses. Human genetic studies consistently associate single nucleotide polymorphisms (SNPs) in the *CHRNA5-A3-B4* cluster, encoding nicotinic acetylcholine receptor (nAChR) subunits α5, α3, and β4, with nicotine addiction by GWAS (Thorgeirsson et al., 2008; Amos et al., 2010; Bierut, 2011). Rare variants in *CHRNA3* have been linked to alcohol dependence in humans (Haller et al., 2014; Kamens et al., 2023). While investigations into contributions of *CHRNA5* and *CHRNB4* to SUD have progressed rapidly using rodent models (Fowler et al., 2011; Frahm et al., 2011), such models for *CHRNA3* are fewer due to neonatal lethality of knock-outs (Xu et al., 1999; Elayouby et al., 2021). This leaves the role of *CHRNA3* in alcohol dependence unclear and correlational. Our study fills this gap, validating *chrna3*’s contribution to the acute alcohol response.

Using SAZA for juvenile zebrafish, we report an acute biphasic response to alcohol, transitioning rapidly from attraction to avoidance. This correlates with the proposed role of alcohol as both stimulant and sedative. *chrna3* mutants showed alcohol tolerance, blunting the avoidance response and resisting changes to shoaling behaviour. In the context of the Genetically Informed Neurobiology of Addiction (Bogdan et al., 2023), our study provides new evidence that variations in *CHRNA3* may directly influence predisposition to alcohol addiction in humans.

## Materials & Methods

### Fish husbandry

Zebrafish (*D. rerio*, ABWT) were housed, bred, and reared as described by Nathan *et al*., 2022 (Nathan et al., 2022) at the ZebraFish Facility (Institute of Molecular and Cell Biology, A*STAR) in groups of 20–30 in 3-L tanks under standard facility conditions. All experimental protocols involving zebrafish were approved by the Institutional Animal Care and Use Committee (IACUC) of the Biological Resource Center at A*STAR. Approved experimental protocols (IACUC #201529, # 231808) were followed.

### Generation of chrna3 mutant zebrafish

The full details of zebrafish gene editing by CRISPR/Cas9 have been described by Hwang *et al. (Hwang et al., 2013)*. CRISPR targets were determined using the web tool ‘chop-chop’ for the exon 5 region to disrupt key functional domains. Oligonucleotides containing the identified CRISPR target sequence (GGN_20_), along with a universal reverse primer encoding the sgRNA scaffold, were used to generate a transcription template, following the method described by Bassett et al. (Bassett et al., 2013). The sequences of these oligonucleotides are listed in Table 1. CRISPR target sites are underlined, and homology arms are shown in lowercase letters. Zebrafish embryos (nacre background) were injected at the 1-cell stage with 1 nL of a mixture containing 1 μL sgRNA (≈ 4-5 μg) and 1 μL Cas9 mRNA (≈1 μg). Fish were genotyped by fin-clipping and sequencing of genomic DNA at 10 weeks post-injection to identify founder fish (F0). The identified founder fish were outcrossed to AB background WT fish to obtain F1 embryos. F1 embryos from each outcrossed family were collected and some of the embryos were verified by sequencing. Once germline transmission was confirmed by sanger sequencing, the remaining embryos were grown to adulthood. Homozygous *chrna3^C246X/C246X^* mutants (AB x nacre mixed background) from the F3 generation onwards were used for all subsequent experiments, alongside WT equivalents, and are henceforth referred to as *chrna3^C246X^*.

### Self-administration for zebrafish assay (SAZA) apparatus setup

The SAZA assay apparatus was set up as described in (Nathan et al., 2022), outlined briefly as follows. The assay tank was constructed of opaque, 3mm thick acrylic to be 35 x 75 x 30mm (width × length × height). One end of the zone contained a dividing sheet of the same material, 30mm in length, to split the zone into two halves while still permitting access (Figure 1A). The zone was placed on an LED lightbox for illumination, providing 5000 lux at maximum intensity (LightPad LX Series, Artograph, USA). Video footage was captured at 30 frames per second using an acA2040-90μM USB3.0 Basler camera. Solutions were dispensed from vertically mounted 10ml syringes through silicon tubing (outer diameter: 1/16”, inner diameter 1/32”) into the corner of their designated zone, mediated via solenoid pinch valves (Automate Scientific, USA, SKU: 02-pp-04i) (Figure 1A). System water was consistently supplied to the tank through additional silicon tubing at the opposite end of the tank to the divided zones and was removed at a matching rate (∼2ml per minute) by gravitational siphoning from the end of the dividing sheet near the centre of the tank (Figure 1A). This created a consistent flow and stopped dispensed solutions from escaping their designated zones while permitting access to the fish being assayed (Nathan et al., 2022). CRITTA (http://www.critta.org), a custom LABVIEW software, was used to track fish movements as described by Krishnan *et al*. (2014) (Krishnan et al., 2014). The tracking system was configured so that only the entry of the fish to a designated region of interest would trigger the pinch valve to release and dispense the corresponding solution.

**Figure 1.**
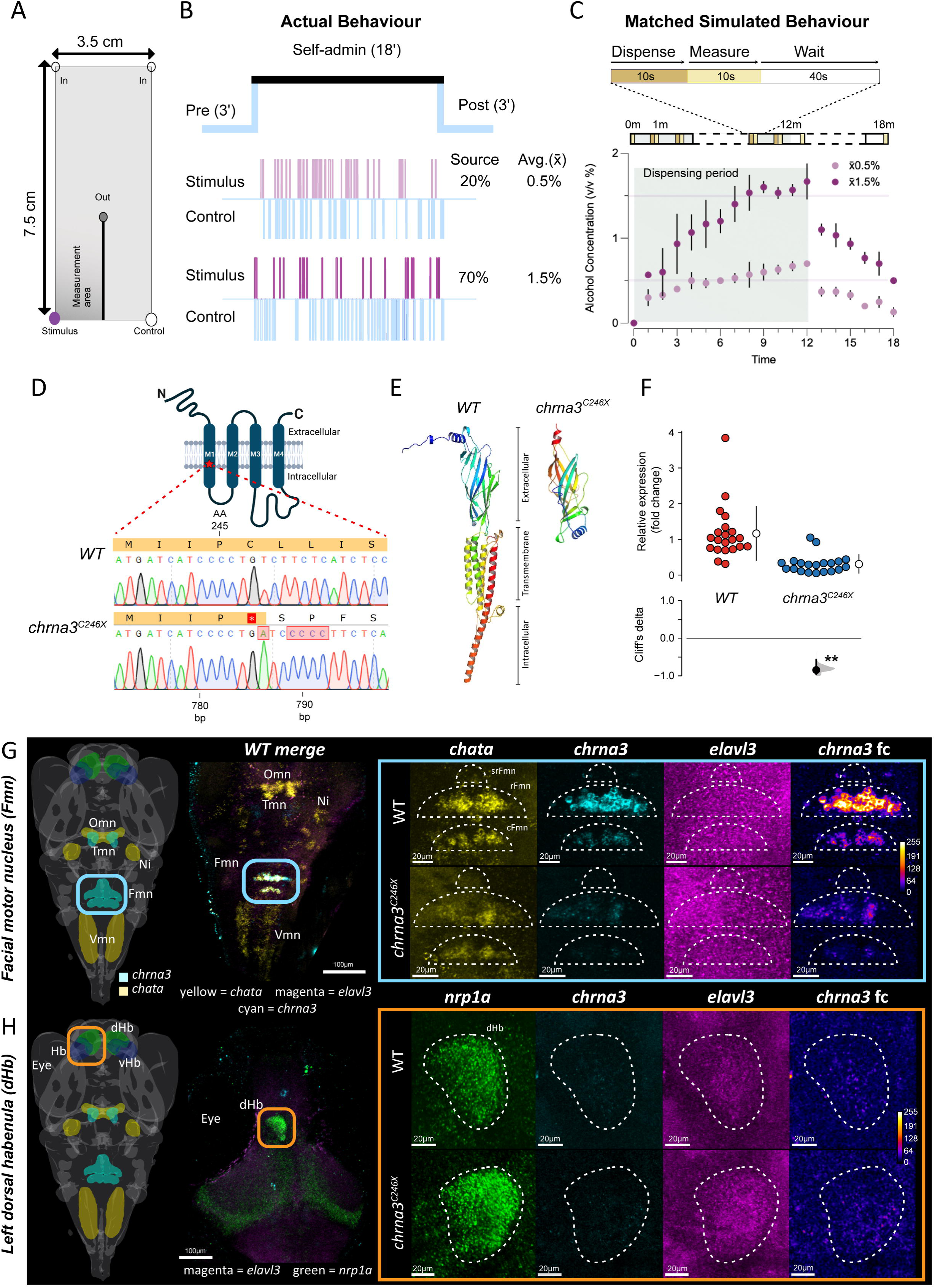
Self-Administration Zebrafish Assay (SAZA) chamber functional design and characterisation of the *chrna3^C246C^* mutant zebrafish. **A.** Schematic indicating locations of inflow of system water (In, top) and outflow (Out, centre). Control and stimulus delivery inlets (stimulus shown in purple) are activated when a fish enters the corresponding zone. Continuous inflow and outflow constrain the stimulus/control delivered to their respective zones (Nathan et al., 2022). ‘Measurement area’ denotes where alcohol concentration was measured in panel C. **B.** Schematic shows the experimental design with three minute pre- and post-exposure periods before and after an 18 minute self-administration period. Representative sparklines plots show the entries and duration (thickness) of entry of the subject fish either into the stimulus or the control zones when 20% or 70% alcohol is used as stimuli. **C.** Alcohol concentrations measured when 20% or 70% alcohol were dispensed for the duration mimicking subject fish behaviour (± standard error). The average concentration (solid line) fish experience is 0.5% and 1.5% alcohol respectively. See Table 2 for additional details. **D-H.** Characterisation of *chrna3^C246C^* mutant fish*. **D***. *chrna3^C246C^* mutants have a premature stop codon in the transmembrane region 1, amino acid 246 (M1 - red star), due to the indel of 5bp at position 786bp. **E.** Predicted protein structure of *chrna3^C246C^* in WT and *chrna3^C246C^* mutants, generated by iTasser (Zhou et al., 2022), colour coded by domain. **F.** Fold change of *chrna3^C246C^* in 12 week old WT (n = 21) vs. *chrna3^C246C^* (n = 20) by qRT-PCR. **G-H.** *chrna3* expression in 14 day old zebrafish larvae visualised by Hybridisation Chain Reaction^TM^ RNA-FISH (HCR^TM^) in the **G.** Predicted facial motor nucleus (Fmn) marked by *chata* expression and, **H.** dorsal habenula (dHb) marked by *nrp1a* expression, in WT and *chrna3^C246C^* respectively. Images are representative, see figures S1 and S2 for replicates (WT/*chrna3^C246C^* Fmn n = 3/4, dHb n = 4/3). Schematic images with regional markers (v2.0, MECE, 2024) were generated in mapZeBrain Atlas (mapzebrain.org, May 2025) (Kunst et al., 2019). Hb = habenula, vHb = ventral habenula, Omn = oculomotor nucleus, Trm = trochlear motor nucleus, Ni = nucleus isthmi, Vmn = vagus motor nucleus, srFmn = supra-rostral Fmn, rFmn = rostral Fmn, cFmn = caudal Fmn. Genes presented left to right: all genes merged, **G.** *chata* or **H.** *nrp1a, chrna3, elavl3*, and *chrna3* with false colour post-processing. False colour adjusted (fc) and elavl3 images have been brightened for enhanced visibility.

### Alcohol concentration measurement

Alcohol concentrations in the SAZA stimulus zone were measured using a DensitoPro (Mettler-Toledo), returning the measured solution to the zone with minimal disturbance. As attempting to measure the concentration of alcohol during a live fish assay would disturb the fish and prevent a natural response, an artificial dispensing scheme of ten continuous seconds per minute followed immediately by measurement was used as a substitute (Figure 1C). This was based on the average time fish spent in the stimulus zone under assorted alcohol dispensing solutions over 18 minutes of exposure (Table 2). After 12 minutes, the dispensing cycle was halted, but measurements continued to estimate the effect the outflow rate had on concentration in the zone if no stimulus was dispensed for a long period. The assay was repeated three times per concentration of alcohol (20, 70%).

### SAZA

SAZA was performed as described in (Nathan et al., 2022), outlined briefly as follows. First, 40 ml of system water was added to the tank, the in and outflow tubes were inserted into their respective conditions, and the flow of system water started. One zebrafish of 30-35 dpf was taken directly from its husbandry tank and added gently to each SAZA tank, then given ∼5 minutes to adjust and recover post-transfer until consistent swimming behaviour had resumed and the fish had explored all zones of the SAZA tank. The recording and tracking scheme was then started for 24 minutes, consisting of three consecutive periods; three minutes of pre-exposure, 18 minutes of self-administration stimulus delivery, and three minutes post-exposure. Only during the 18-minute stimulus delivery period would entry to a designated zone trigger dispensing of the corresponding solution (Figure 1A). The valve opening time was set to a minimum of five seconds per entry. The stimuli delivered were 0/20/70% absolute alcohol diluted in the facility system water. Following each assay, the zone was emptied of all liquid, washed, and refilled before adding the next fish. The volume of each solution dispensed was also recorded. To reduce bias, the zone designated to deliver the stimulus solution was randomly assigned to be either the left or right and changed periodically throughout the experiment such that ∼50% of the fish were tested in each configuration. Around 30 fish naive to SAZA were assayed per concentration of alcohol. Following each assay, the fish were euthanized. Juvenile fish, ∼5 weeks old, were used for all behavioural experiments as this developmental stage offers the advantages of more complex locomotor behaviour and reduced individual variability over younger larvae, while possessing comparable structural brain development to adult fish, yet still facilitating higher throughput and practicality of assay design.

### SAZA Data analysis

SAZA data outputs from CRITTA were processed by custom Python scripts. These provided data on time spent in each zone and mean velocity in each zone used in these analyses. The data were separated into three major time groups; pre-exposure, total stimulus delivery period, and post-exposure. The stimulus delivery period was then further subdivided into either start and end (0-9m/9-18m), or three minute segments (0-3m/3-6m/6-9m, etc…) for further analyses.

The preference index (PI) was calculated as the ratio of the difference between the times spent in the stimulus and control zones relative to the total time spent in the two zones. The PI for a given duration (PI_X_) was calculated by the following formula; PI_X_ = (TinS - TinC) / (TinS + TinC), where TinS = time in stimulus zone (s) and TinC = time in control zone (s). This gives the value for PI maximum bounds of + or - 1 indicating more time spent relatively in the stimulus or control, respectively. For example, a PI of +1 would indicate 100% of relative time spent in the stimulus zone during one time period. Delta PI between two time periods (ΔPI_X-Y_) was calculated as ΔPI_XY_ = PI_Y_ - PI_X_, where PI_X_ = the earlier period, and PI_Y_ = the later period. This gives the value for ΔPI maximum bounds of + or - 2, with the values indicating the direction and magnitude of the change in preference from the first time point to the second. For example, a ΔPI of +2 would indicate a change from 100% relative time spent in the control zone in the first period, to 100% relative time spent in the stimulus zone in the second. Sparklines were also generated to display the number of entries to each zone, and the duration of each (Figure 1A).

### Shoaling cohesion assay

The shoaling cohesion assay apparatus was set up as described for SAZA, with the following alterations. The tank had no dividing sheet, or inflow, outflow, and stimulus dispensing tubing (Figure 4A). Additionally, the tracking of fish movements was not performed. To perform the assay, the tank was half-filled with 20 ml of system water, then four zebrafish of 30-35 dpf were taken directly from their husbandry tank and added gently to each shoaling cohesion tank. The fish were then given ∼5 minutes to adjust and recover post transfer until consistent swimming behaviour had resumed. Following this acclimation period, 20 ml of 2x concentration alcohol solution was added to the tank to achieve the 1x desired concentration for the assay. Recording began immediately and continued for 30 minutes. Six repeats were performed per alcohol concentration. Following each assay, the fish were euthanized.

### Shoaling cohesion data analysis

Video analysis was performed in imageJ. First, videos were either trimmed to the first 15 minutes, then frames were extracted at 2fps to create an image stack (900 per assay), or one frame per 10 seconds of footage was extracted from the full 30-minute duration (180 per assay). The image stacks for each assay were cleaned using median background subtraction, then converted to 8-bit and auto-thresholded (Figure 4A). From the thresholded images, particles were detected (size = 100-infinity), and the centerpoint of each was determined. Using the centerpoints, Delaunay triangulation was performed on each image to draw triangles covering the space between all four fish. The area of these triangles was then combined to obtain the total area between the fish, as a measure of shoaling cohesion. These area values were grouped into either total time for the full 30 minute duration (n = 1080 per treatment), or one minute intervals for the first 15 minute data (n = 720 per time point per treatment). Values over 2000 mm^2^ were identified as products of processing errors and removed (∼2.5%).

As a measure of shoal kinesis, the distance travelled by the shoal centerpoint was calculated using the X and Y coordinates for the centerpoint of the triangulation area, as calculated by imageJ, with the equation; distance = √((X_2_ – X_1_)² + (Y_2_ – Y_1_)²). X_2_ and Y_2_ were the subsequent coordinates, 0.5s after X_1_ and Y_1_, respectively. This provided the distance the shoal centerpoint had travelled between each selected frame. Values over 10 mm per 0.5s were discarded as products of processing errors, as the highest velocities recorded by individual fish were ∼20mm/second (Figure 3B). This amounted to < 5% of the data. These values were grouped into one minute intervals for the first 15 minute data (n = 720 per time point per treatment).

### Quantitative PCR (qPCR)

Twelve-week old adult zebrafish were euthanized and the brains were dissected in PBS pH7.0. Brain tissue was homogenised using a micro tube homogenizer (Thomas Scientific) in a lysis buffer made up of Trizol (ThermoFisher #15596026). RNA was extracted using PureLink® Micro Kit (ThermoFisher #K310250) according to the manufacturer’s protocol. The concentration and purity of the extracted RNA were determined by NanoDrop™2000 Spectrophotometer (ThermoFisher). Each brain sample yielded around 100 ng/μL RNA with a ratio of absorbance reading 1.9-2.0 at 260/280 nm. The purified RNA was reverse transcribed by SuperScript II First strand Synthesis System (Invitrogen #18091050) using 100 ng/μL of purified RNA to obtain approximately 1000 ng/μL of first strand cDNA. To check for genomic DNA contamination, a negative RT (no reverse transcriptase) was set up for each sample.

The cDNA was diluted with nuclease-free water to 100 ng/μL. The qRT-PCR amplification mixtures (20 μL) contained 100 ng of cDNA, 10 μL 2x GoTaq®qPCR Master Mix (Promega #TM318), and 300 nM forward and reverse primer. The primers against *chrn* targets used were designed using Primer3 (v.0.4.0) and are detailed in Table 3.

Reactions were performed using a 7500 Real-Time PCR system (Applied Biosystems) in 96-well plate format. The cycling conditions were as follows: 95°C for 10m to activate the polymerase, then 40 cycles of 95°C for 15s, 60°C for 30s. After completion, a melt curve was run at 65-95°C for 5 seconds per step. All PCR efficiencies were above 95%. Primer specificity was validated by a single peak on the post-PCR melt curve and single-band after electrophoresis.

The relative gene expression levels were analysed using the adjusted delta-delta CT method described by Hellemans and Mortier et al. (2007). All data were analysed by one-way ANOVA, with genotype as the independent variable and target gene expression, relative to reference genes, as the dependent variable. For mRNA relative expression data, adjusted *p* values were estimated using Dunnett post-hoc tests. Effect sizes for all differences in expression were also calculated using Cliff’s Delta statistics as described below.

### Statistical analyses

Data was analysed and presented following the principles outlined by Ho *et al*. (2019), to which a p value as a measure of significance is improved upon by the supplementary reporting of effect sizes, means, and 95% confidence intervals (Ho et al., 2019). As the assumptions of normality and homoscedasticity recommended for parametric testing could not be consistently met for the SAZA data, most unpaired data were compared by non-parametric two-sided permutation T-tests, (5000 reshuffles) with a significance level of 0.05, and unpaired Cliff’s delta, while paired or repeated measures data were compared using by paired permutation tests under the same parameters. Due to the more complex design and higher replicate availability, the shoaling assay longitudinal data were analysed using a linear mixed effects model, with shoal area or centerpoint travel distance as the response variable, alcohol treatment and time as fixed effects, and assay ID as a random effect to account for repeated measures. The residuals indicated the data were slightly skewed but appeared normally distributed and homoscedastic enough to support a parametric test. Analysis was performed in R using the ‘dabestr’ (v2024.12.24) and ‘lmerTest’ (v3.1-3) packages. In the case of multiple comparisons, the Holm-Bonferroni step down family wise error correction was applied to the *p* value. All statistical analysis outputs can be found in supplementary tables. This data was presented in the form of Gardener-Altman and Cummings estimation plots. Results are reported as Cliff’s delta, or Paired mean difference = X, 95% CI [lower, upper], *p* = Y. For Cliff’s delta, effect sizes greater than ± 0.4 and a *p*-value of < 0.01 was considered meaningful and practically relevant in this study. Smaller Cliff’s delta effect sizes of ± 0.2 to 0.4 with accompanying *p*-value of < 0.05 were also considered potentially meaningful, but provisional pending experimental reproduction (Cliff, 1993; Goodman et al., 2019).

### Hybridisation Chain Reaction, RNA-fluorescent in-situ hybridisation (HCR^TM^ RNA-FISH)

Hybridisation Chain Reaction RNA-FISH 3.0 (HCR) (Choi et al. 2018) probes against *chrna3*, *chata*, and *nrp1a* were prepared using an in-house Excel-based program and synthesized by IDT (please see Table 4 for details). The in-situ hybridization experiments were carried out in a controlled incubation chamber (40°C temperature and > 90% humidity). Briefly, 14 dpf zebrafish embryos were euthanized and fixed in 4% formaldehyde diluted in 1X PBST at room temperature for 60 minutes with shaking. The larvae were washed with 1X PBST for 3 minutes x 3 times. Embryos were permeabilized using a detergent solution (1% SDS, 0.5% Tween 20, 150mM NaCl, 1mM EDTA, 0.05M Tris-HCL, pH 7.5) at 37°C for 30 minutes. They were washed for 3 minutes x 3 times each with 1X PBST and then 5X SSCT. Larvae were transferred to 30% probe hybridization buffer (30% formamide, 9mM citric acid, 0.1% Tween 20, 50µg/mL heparin, 1x Denhardt’s solution, 5X SSC) and incubated at 37°C for 30 minutes. Simultaneously, probes were prepared in a solution containing 30ul (10uM) of probe set against each gene in 1000 µl of 30% probe hybridization buffer. The larvae were either transferred to the robotic chamber or a 1.5ml centrifuge tube for automated or manual experiments. Larvae were incubated with the probes at 37°C for 8-12 hrs. For both methods, on day 2, larvae were washed 5 times with 30% probe wash buffer (30% formamide, 9mM citric acid, 0.1% Tween 20, 50µg/mL heparin, 5x SSC) at 37°C and then 5X SSCT twice. For the secondary reaction, HCR hairpins (Molecular Instruments) were diluted and prepared in an amplification buffer (5X SSC, 0.1% Tween 20, 2.5% dextran sulphate) and incubated overnight (for the manual methodology) or 3 hours (for the automated) in the dark. The larvae were then washed with 5X SSCT, 4 X 10 mins each in the dark. They were mounted with the cover slip on the dorsal side in 70% glycerol-PBS mounting media and imaged on an Olympus FV3000 confocal microscope with the settings consistent between fish. Images were post-processed first on Olympus viewer FLUOVIEW FV31S-SW software with linear intensity adjustments of 150 - 4095 for all genes. Images presented for *nrp1a*, *chata* and *chrna3* had no further processing beyond cropping to the ROI and addition of scale bars. The *elavl3* and *chrna3* false colour images were processed in Imagej. The *elavl3* images had additional linear brightness/contrast adjustment (15 - 110). False colour images were converted to 8bit and had the ‘fire’ LUT applied, followed by linear brightness/contrast adjustment of 0 - 175, background subtraction with 25px rolling ball, and 1px gaussian blur. Fluorescence quantification (S1, S2) was performed in ImageJ on the raw images without post-processing. The regions of interest were marked using the boundaries observed in *chata* for the facial motor nucleus, or *nrp1a* for the dorsal habenula, and mean grey pixel value within the region was measured. Two week old (14 dpf) larvae were used for HCR imaging as this age represented the upper limits of transparency without additional optical clearing steps, while being closer to behavioural testing age than the traditional imaging of 5-7 dpf larvae. Single cell analysis shows that 97% of the cell types in a 15 dpf fish are present in 25 dpf fish (Raj et al., 2020), so we deemed it an acceptable compromise for the benefits throughput and visualisation over imaging older animals.

### RNAseq

RNA was extracted from twelve-week old adult zebrafish *chrna3* and WT brains in triplicate as described previously. The quality of each sample was verified by gel electrophoresis and bio-analyser. Samples that passed QC were sent for sequencing by Novogene AIT. An mRNA cDNA library made up of Paired-end sequencing reads of 150 bp in length was generated on the Illumina HiSeq instrument. Raw reads were first processed by Novogene-AIT to remove adapter, poly-N sequences, and reads with low quality from raw data. All downstream analyses were based on the clean data with a Phred score of 39 indicating a 99.9% base call accuracy. Further analysis was performed in Partek™ Flow™ Explore Spatial Multiomics Data using Partek™ Flow™ software, v11.0. Reads were aligned to GRCz11 genome assembly by STAR aligner, default parameters, and subsequently filtered to a minimum mapping quality score of > 30. Gene counts were normalised by median ratio, followed by differential expression analysis in DeSeq2. Significance thresholds were set as follows: up/down-regulated genes = p < 0.05 & fold change > ±2. Provisional up/down-regulated genes = p < 0.05 & fold change < ±2. Non-significant genes = p > 0.05. Custom gene lists and sets for analysis can be found in Table 5. Gene set enrichment analysis (GSEA) was performed using the GRCz11-GO database, 100 permutations. Significance thresholds were set at p < 0.05 and FDR < 0.2. Given the loss of spatial and structural resolution when using bulk RNA sequencing, this investigation was primarily targeted at identifying brain-wide, compensatory mechanisms that could potentially underlie phenotypic changes, or seed predisposition to multi-substance dependence and comorbid disorders, we opted for the ‘fully developed’ brain of the sexually mature adult. These larger adult tissues provided larger quantities of high quality RNA more consistently than younger animals.

## Results

### Zebrafish preference for alcohol self-administration in SAZA is biphasic

We refined the Self-Administration Zebrafish Assay (SAZA) first reported in (Nathan et al., 2022) to improve the temporal resolution for quantifying the dynamic nature of the acute response. SAZA apparatus created three chemically distinct, but physically continuous zones, or regions within the assay chamber. The inflow and outflow were adjusted by gravity feed to maintain the integrity of the two delivery zones throughout the assay period (Nathan et al., 2022). Subject fish behaviour was monitored by a closed-loop, video acquisition program developed in-house that allowed for self-administration of a stimulus or a control when subject fish entered a delivery zone from a neutral zone (Figure 1A; Movie 1). To avoid intrinsically preferred swimming direction bias, the stimulus and control delivery zones were pseudo-randomly assigned in each trial such that half of the fish in each group received the stimulus (alcohol) in the left and the other half in the right delivery zone (Figure 1A). Sparkline plots in pilot experiments showed that fish administered 20%, or 70% alcohol (at source) differentially, by changing the frequency of entry, and duration of time spent in the stimulus area (Figure 1B; Movie 1). This suggested that fish could quantifiably self-regulate alcohol administration, and their behaviour in the closed-loop setup was dependent on stimulus concentration.

Next, we estimated the stimulus concentration that fish experienced during the assay. To do so, we used a densitometer to measure the change in water density caused by the alcohol-water mixture. Alcohol was dispensed mimicking the behaviour of fish in the pilot experiment (Figure 1B, ∼10 seconds/min discontinuously, table 2). This analysis revealed that, due to improved outflow and the lower density of the alcohol-water mixture, the alcohol concentration in the chamber was only a fraction of that at the source. This concentration was lower than previous estimates obtained from colorimetric analysis of food dye-containing solution (Figure 1C). The maximum concentration plateaued at ∼40-fold lower than the source alcohol concentration, which was at least four-fold lower than previous studies (Nathan et al., 2022). Therefore, on an average, self-administering source alcohol at 20% and 70% resulted in the fish experiencing alcohol at ∼0.5% and ∼1.5% respectively in the assay (Figure 1C; for 20%, 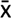 = 0.52 %, SD = 0.13%; 70%, 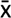 = 1.52%, SD = 0.44%). We refer to these two conditons as x0.5% conditon and 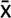1.5% respectively in the rest of this study. The effective concentration reduced by half within a minute of stopping self-administration (Figure 1C). Thus, both the pre, and post-administration periods served as controls to compare the behaviour of animals in the naïve pre, and post exposure periods. This range of alcohol exposure is comparable to the concentration of alcohol used in previous zebrafish studies (Kurta and Palestis, 2010; Echevarria et al., 2011; Mathur et al., 2011; Tsang et al., 2019).

The dynamic change in alcohol concentration over time during SAZA is a key advantage for untangling the small effects single genes may have on complex behaviours like alcohol preference. Common variants in *CHRNA3*, for example, are risk factors for developing nicotine dependence, while several rare variants are proposed to influence cocaine and alcohol dependence (Haller et al., 2014; Kamens et al., 2023). *Chrna3* is proposed to mediate alcohol stimulant effects in rodents (Kamens et al., 2009). However, this gene’s role in development of alcohol dependence hasn’t been tested directly. We hypothesised that the SAZA will be effective in examining such a role, if present, in nicotinic acetylcholine receptor alpha3 (*chrna3)* mutant generated recently (Kibat, 2020).

The *chrna3* mutant harbours an insertion that causes a frameshift and introduces a premature stop codon within transmembrane domain 1 of the gene, resulting in a truncated protein at amino acid position 246 (*chrna3*; Figure 1D). The mutant protein is predicted to have a large truncation of the transmembrane and the intracellular domains (Figure 1E) and is thus expected to be damaging to protein function. Quantitative RT-PCR of adult whole brains showed that the mutation resulted in a significant reduction in the expression of *chrna3* transcripts in the mutants (Figure 1F, *WT* vs. *chrna3*, Cliff’s delta = −0.848, 95CI [−0.567, −0.962], *p* = < 0.001). In rodents, *Chrna3* expression is restricted. It is expressed in a few neurons in the hippocampus, medial habenula, interpeduncular nucleus (IPN), the ventral tegmental area (VTA), cerebellum, locus coeruleus, and spinal cord (Wada et al., 1989; Gotti et al., 2006). We investigated the expression of *chrna3* in 14 days post fertilization (dpf) zebrafish larvae using an in-house modified Hybridisation Chain Reaction™ RNA fluorescence in-situ hybridisation (HCR) method (Banerjee et al., 2024). As cholinergic function in the hindbrain motor neurons and expression in facial motor nucleus (Fmn) have also been previously reported (Rima et al., 2020), we quantified *chrna3* expression in the mutants. While expression of *chrna3* was high in the wild types, it was starkly and consistently reduced in the mutants (Figure 1G, S1). We also examined expression of *chrna3* in brain regions such as the medial habenula (dorsal habenula in zebrafish), reported to have co-expression of *CHRNA5-A3-B4* in mammals (Xu et al., 2006; Fowler et al., 2011). Only a few cells in the left dorsal habenula expressed *chrna3* at a low level. The expression levels in the mutant and WT were similarly low (Figure 1H, S2).

Having established a robust system for quantifying self-administration and generated *chrna3* mutants, we examined the behaviour of juvenile fish in the SAZA at an age when they show consistency across the population (∼5 weeks old). We first examined the response of wild-type animals (WT). The pilot (Figure 1B) suggested the preference changed dynamically over the course of the assay. To capture this, we collapsed the preference behaviour of animals over 3-minute windows that balanced temporal granularity in relation to the changing alcohol concentration with quantifiable observation periods. The preference was unchanged over time when alcohol was absent (0%; Figure S3). The same analysis for the 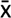0.5% alcohol condition revealed that fish initially spent significantly more time in the stimulus zone compared to the pre-exposure period (Figure 2A, 3-6m, Paired mean difference = 7.62, 95CI [13.9, 0.331], *p_adj_* = 0.028). However, after about 10 minutes, fish reversed their preference and spent significantly less time in the stimulus zone (Figure 2A, 12-15m, Paired mean difference = −6.47, 95CI [−2.2, −10.1], *p_adj_* = 0.031). This avoidance continued for the remainder of the assay, even after the self-administration period ended. However, the early attractive portion of this biphasic response was absent in the 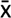1.5% condition (Figure 2B, 3-6m, Paired mean difference = −1.1, 95CI [4.26, −6.27], *p_adj_* = 1). Fish spent less time in the stimulus zone from six minutes onwards (Figure 2B, 6-9m, Paired mean difference = –8.57, 95CI [−3.22, −13.00], *p* = 0.012). Mapping the behavioural changes to the alcohol concentration expected in the stimulus zone revealed that juvenile zebrafish preferred alcohol when the concentration was under 0.5%, while they avoided ∼0.7% or higher alcohol (Figure 2C). Thus, WT zebrafish self-regulated alcohol exposure within a narrow range.

**Figure 2.**
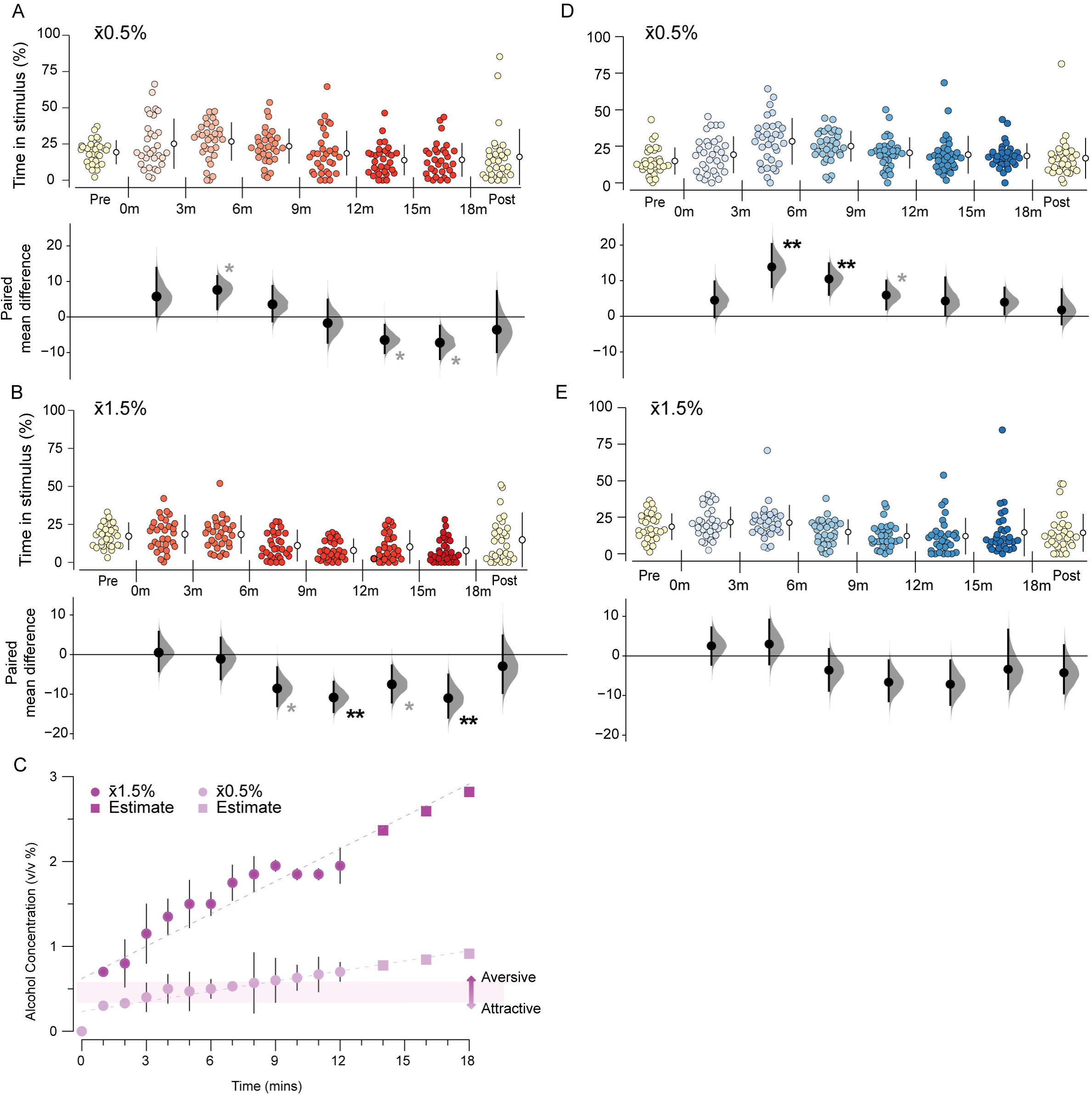
The preference for alcohol is biphasic. Gardener-Altman and Cumming estimation plots show alcohol response in three-minute intervals. No stimulus was dispensed in the ‘Pre’ and ‘Post’ periods. Time spent in the stimulus zone during 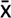0.5% SAZA condition for **A.** WT (n = 31) and **D.** *chrna3^C246C^* mutants (n = 32). Time spent in the stimulus zone in the 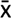1.5% SAZA condition for **B.** WT (n = 30) and for **E.** *chrna3^C246C^* mutants (n = 30). **C.** Ethanol concentrations in the stimulus zone in the 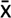0.5% and 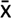1.5% alcohol conditions and estimated for extended periods. The concentration at which attraction switches to aversion is highlighted. Asterisks indicate a significant paired mean difference between that time portion and the pre-stimulus period: * = p_adj_ < 0.05 (provisional difference), ** = p_adj_ < 0.01 (meaningful difference). See Table S1 for the precise effect sizes and p-values, corrected for multiple comparisons.

### *chrna3* mutant fish show higher tolerance to alcohol

We next examined the behaviour of *chrna3* mutants in the SAZA. Mutants, similar to WT counterparts showed no preference in the absence of alcohol (0%; Figure S3, A-B). In the 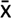0.5% condition, *chrna3* fish initially behaved like their WT spending significantly more time in the stimulus zone (Figure 2D, 3-6m, Paired mean difference = 13.9, 95CI [20.3, 8.18], *p_adj_* = 0.005). However, unlike the WT, this attraction persisted throughout the assay period (Figure 2D, 9-12m, Paired mean difference = 5.96, 95CI [10.1, 1.94], *p_adj_* = 0.044). No biphasic switch to avoidance was observed. Surprisingly, mutants did not show a strong avoidance response during the assay even in the 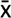1.5% condition (Figure 2E, 12-15m, Paired mean difference = −7.15, 95CI [−1.13, −12.4], *p_adj_* = 0.134).

The total time spent in the control delivery zone varied among conditions and genotype, but did not show a consistent trend (Figure S3, C-H). The frequency of entry into the stimulus zone however, was correlated with the time spent in the stimulus zone. This was similar across both genotypes in the 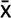0.5% condition and was indicative of the preference behaviour seen (Figure S4, C-D). Interestingly, the frequency of entry into the stimulus zone in the 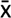1.5% condition showed a brief increase initially across the two genotypes (0-3 mins; Figure S4, E-F), before it plateaued off to pre-stimulus frequency rapidly, suggesting that a reduction in total time spent in the stimulus zone observed in Figure 2B and 2E was driven by shorter visits, rather than reduced frequency of visits.

Despite the early attraction (Figure 2A, 2D), in general alcohol appeared to be aversive to WT zebrafish albeit less so for the mutants. This could also be inferred from the volumes delivered in alcohol conditions compared to the volumes delivered in the 0% alcohol condition (Figure 3A, 3B). WT fish self-administered significantly smaller volumes in the 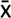0.5% condition (Figure 3A, Cliff’s delta = −0.554, 95CI [−0.266, −0.759], *p_adj_* = <0.001), and in the 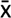1.5% condition (Figure 3A, Cliff’s delta = −0.728, 95CI [−0.441, −0.895] *p_adj_* = <0.001). This supports previous observations that alcohol can be aversive to zebrafish at certain concentrations (Nathan et al., 2022). The *chrna3* mutant fish however, reduced volume of administration only in the 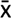1.5% condition (Figure 3B, 0% vs 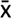1.5%, Cliff’s delta = −0.583, 95CI [−0.304, - 0.791], *p_adj_* = < 0.001), and the two genotypes differed only during the 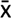0.5% condition (Figure 3C, 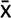0.5%, Cliff’s delta = −0.328, 95CI [0.584, 0.009], *p_adj_* = 0.027).

**Figure 3.**
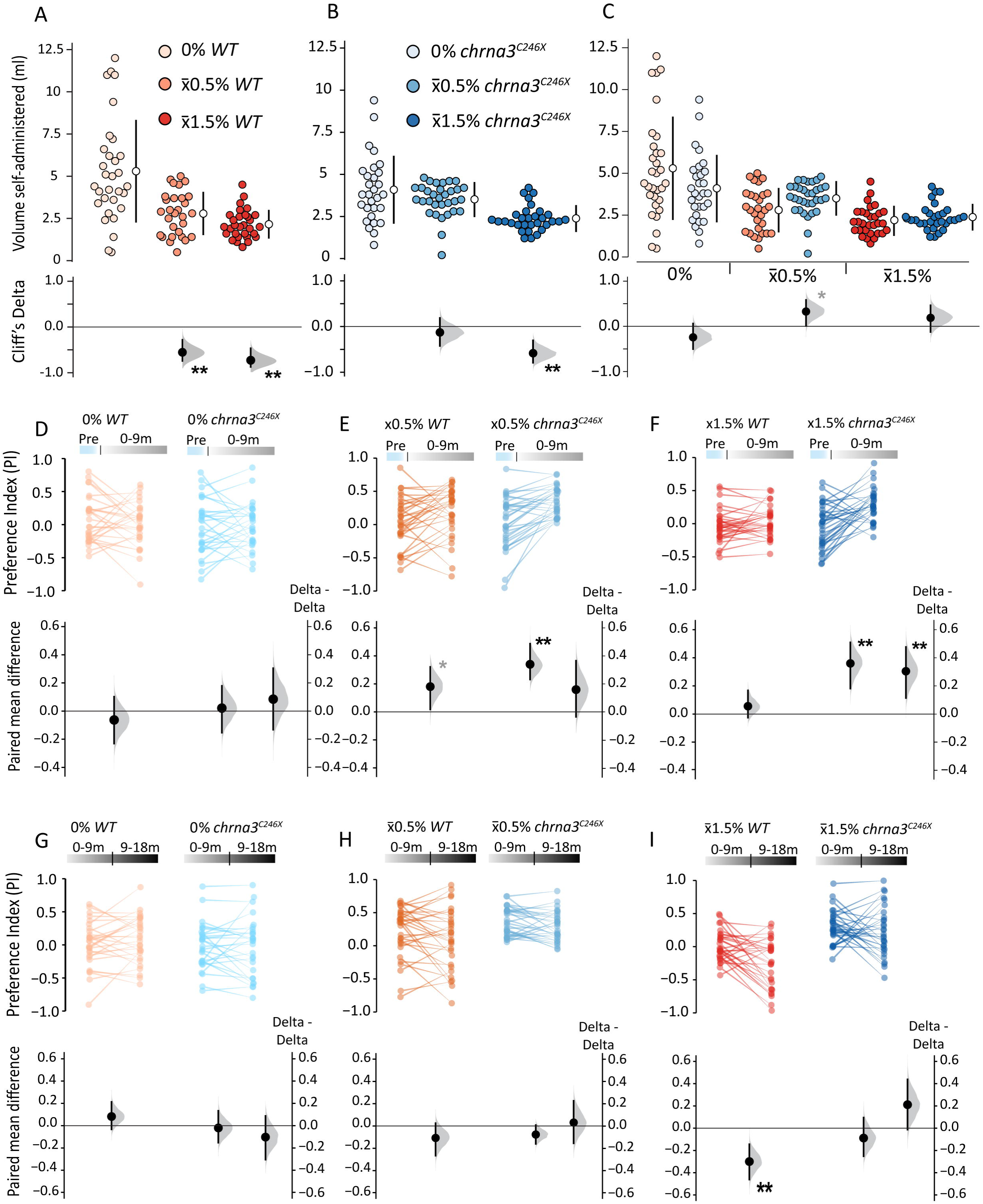
*chrna3* mutation dynamically alters relative preference for alcohol. **A-C.** Gardener-Altman and Cumming estimation plots comparing the volume of alcohol dispensed over the entire self-administration period during SAZA for **A.** WT (0%/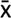0.5%/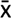1.5%, n = 30/31/30), **B.** *chrna3^C246C^* (0%/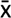0.5%/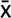1.5%, n = 29/32/30) and **C.** intergenotype comparisons. Asterisks indicate the following significant differences from 0% alcohol treatments (**A, B**), or the WT (**C**): * = p_adj_ < 0.05, and effect size reported by Cliff’s delta between ± 0.2 and ± 0.4 (provisional difference), ** = p_adj_ < 0.01 and effect size bigger than ± 0.4 (meaningful difference). **D-I**. Paired mean difference of preference index (PI) between **D-F**. Pre and 0-9m or **G-I**. 0-9 and 9-18m time periods within genotypes, and delta delta comparisons between genotypes, during **D, G**. 0%, **E, H**. 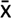0.5%, and **F, I**. 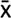1.5% SAZA. Asterisks indicate a significant paired mean difference, or delta delta, between time periods: * = p < 0.05 (provisional difference), ** = p < 0.01 (meaningful difference). See Table S2 for the exact effect size and p-values.

To quantify the magnitude of the difference between genotypes we examined a Preference Index (PI; see methods). PI was calculated for each subject as the relative preference for the stimulus compared to control. Positive PI values (0 to +1) therefore indicated a preference for stimulus, while negative values (0 to −1) indicated an aversion. We also quantified a delta, or the change in the PI itself in subsequent phases: pre-exposure phase, the early phase (0-9 minutes), and the late phase (9-18 minutes). The duration of the phases coincides with the time required (∼10 minutes) for equilibration of blood alcohol concentration in larvae immersed in alcohol (Lockwood et al., 2004). We used the “delta-delta”, or the difference of this measure between the two genotypes to quantify what aspect *chrna3* contributes to most (Figure 3).

As anticipated, the PI range was broad, and population mean was centred around zero in the 0% alcohol condition (Figure 3D). No significant change was observed between pre and early phases for either genotypes. Consequently, the delta-delta between genotypes was also insignificant (Figure 3D). However, both WT and mutants showed a positve preference shift in the early phase in the 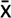0.5% condition (Figure 3E, WT, Paired mean difference = 0.181, 95CI [0.321, 0.017], *p_adj_* = 0.036, Figure 3E, *chrna3*, Paired mean difference = 0.34, 95CI [0.488, 0.232], *p_adj_*= < 0.001). The delta-delta between genotypes changed only marginally (Figure 3E). The genotypic however, was significantly different in the 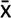1.5% SAZA (Figure 3F, Delta-Delta = 0.305, 95CI [0.478, 0.114], *p_adj_*= 0.009). In the second half of the SAZA self-administraton period, once again the 0% conditon had no directonal change between the phases, or the genotypes (Figure 3G). Although both genotypes showed similar trend in the 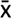0.5% condition (Figure 3H), it was notable that the individual variance in the PI was small for mutants, while it was considerably broader for WT, distributed across preference and aversion. Finally, in the 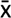1.5% condition, the PI shifted towards aversion, or negative values substantially for WT (Figure 3I, WT, Paired mean difference = −0.3, 95CI [−0.144, −0.462], *p_adj_* = 0.001). One the other hand, the PI in the mutants showed no directionality (Figure 3I). In addition to this increased alcohol tolerance in the 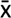1.5% condition the mutants also exhibited heightened alcohol-induced locomotor stimulation over the full stimulus-dispensing period (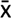0.5% condition; Figure S5, A-F) that impacted other behavioural characteristics of the mutants (Figure S6). Altogether, this suggested that manipulation of *chrna3* function exerted larger effects on the alcohol tolerance at higher alcohol concentrations than impact the attractive response at lower concentrations.

### Shoaling behaviour shows biphasic effects of alcohol exposure

We next explored the potential reasons for this rapid and biphasic response to alcohol self-administration. Alcohol is well-documented to act as an anxiolytic, reducing a variety of anxiety-like behaviours in fish (Gebauer et al., 2011; Araujo-Silva et al., 2020). However, it is also known to function as a sedative, causing reduced swimming velocity and coordination, and increased immobility (Echevarria et al., 2011; Tsang et al., 2019). We hypothesised that the biphasic response in SAZA is correlated to the concentrations at which these two contrasting effects of alcohol come into play. To test this, we examined the shoaling behaviour of juvenile fish. Shoal members are proposed to swim near others when anxious and are expected to be more exploratory when the anxiety-like state is reduced (Miller et al., 2013), for example under an anxiolytic alcohol dose. This will be reflected by an increase in the total area occupied by the shoal, or decreased shoal cohesion. On the other hand, shoal members experiencing the sedative effects of alcohol are expected to swim with decreased speed and coordination. This impacts the distance travelled by the shoal, or the shoal kinesis, in addition to reducing the shoal cohesion. Hence, the impact of both the anxiolytic and sedative effects of alcohol can be measured by examining shoal cohesion and kinesis in tandem (Figure 4A). Shoal cohesion was quantified by defining each fish as a point, then triangulating the lines between them to give the total area between fish. Shoal kinesis was calculated by the distance between the shoal centre points, as defined by the crossing point of the triangulation between fish, between frames at 2fps (Figure 4A).

**Figure 4.**
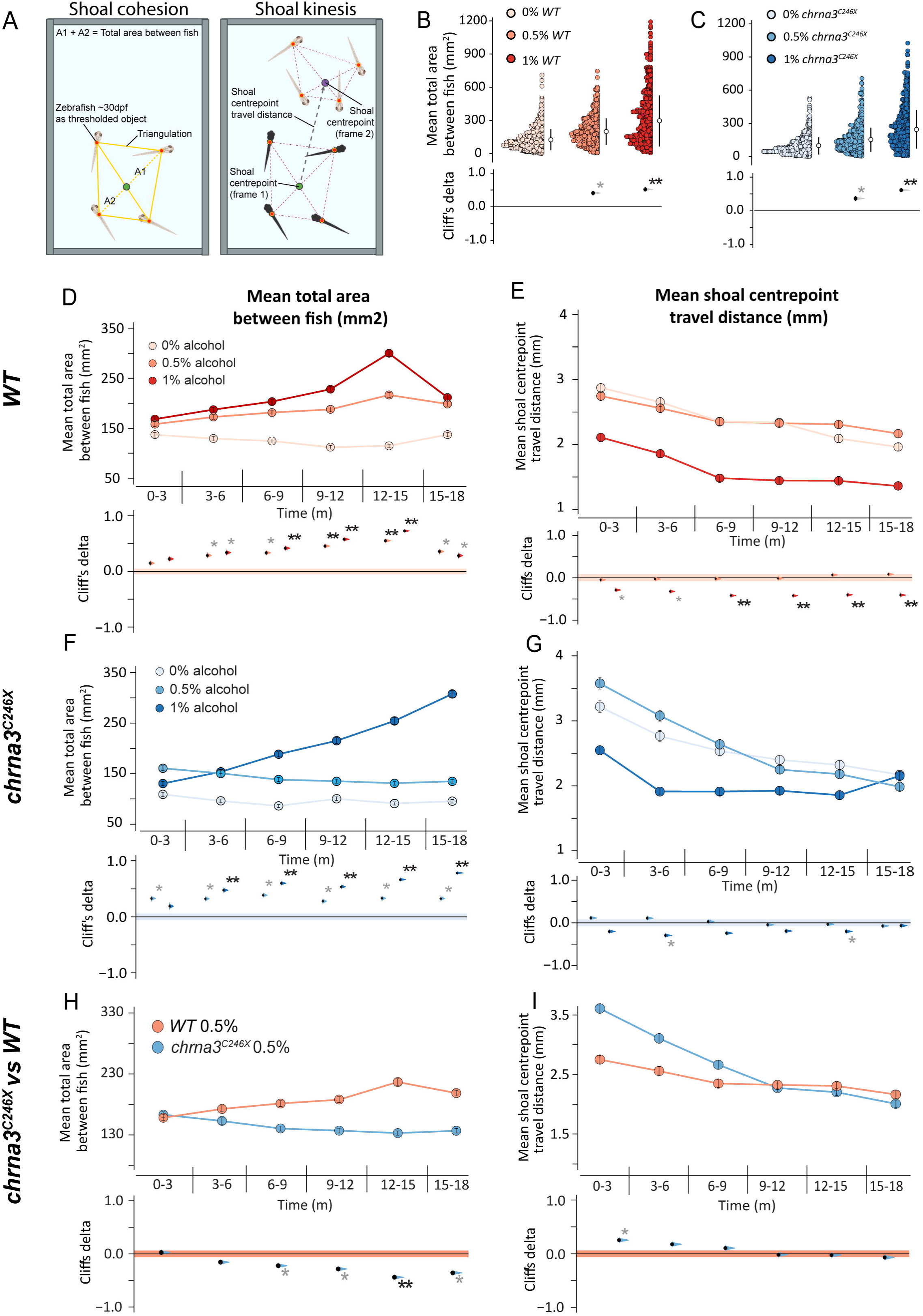
*chrna3* mutants experience reduced psychoactive effects from alcohol exposure. **A.** Schematic outline of the shoaling assay metric calculation and processing steps for cohesion (shoal area) and kinesis (shoal movement). **B, C.** Gardener-Altman, or **D, F, H.** mean (± 95% CI) shoal cohesion, and **E, G, I.** kinesis, with Cumming estimation plots for the shoaling cohesion assay showing **B, D, E.** WT, **C, F, G***. chrna3^C246C^*, and **H, I.** intergenotype comparisons. Metrics recorded per frame at 2fps (n = 6 assays per genotype, per treatment). Asterisks indicate a significant difference in the estimated marginal mean from **B-G.** 0% alcohol, or **H, I.** WT within that time period: * = p_adj_ < 0.05, and effect size reported by Cliff’s delta between ± 0.2 and ± 0.4 (provisional difference), ** = p_adj_ < 0.01 and effect size greater than ± 0.4 (meaningful difference). See Table S3 for exact effect size and p-values.

Taking into account the concentrations associated with attraction and aversion (Figure 2C), we examined the shoaling behaviour of four fish over 30 minutes at two fixed concentrations of 0.5%, or 1% alcohol and compared their behaviour when alcohol was absent (0%; Figure 4A, Movie 2). The total area between WT shoal members increased at both concentrations, with the 1% alcohol having a proportionally larger effect on the shoal area (Figure 4B, 0 vs 0.5%, Cliff’s delta = 0.409, 95CI [0.453, 0.364], *p_adj_* = < 0.001, 0 vs 1%, Cliff’s delta = 0.518, 95CI [0.559, 0.475] *p_adj_* = < 0.001). The same phenomenon was also observable in the mutants (Figure 4C, 0 vs 0.5%, Cliff’s delta = 0.365, 95CI [0.412, 0.320], *p_adj_* = < 0.001, 1%, Cliff’s delta = 0.611, 95CI [0.649, 0.573] *p_adj_*= < 0.001).

Differences between the two genotypes emerged when their behaviours were examined at a higher temporal resolution. Loss of cohesion in WT fish was observable within five minutes of exposure to either concentration (Figure 4D, 0 vs 0.5% 3-6m, Cliff’s delta = 0.287, 95CI [0.320, 0.252], *p_adj_* = 0.032; Figure 4D 0 vs 1% 3-6m, Cliff’s delta = 0.338, 95CI [0.370, 0.305], *p_adj_* = 0.004). The magnitude of the difference increased over time (Figure 4D, 0 vs 0.5% 12-15m, Cliff’s delta = 0.552, 95CI [0.579, 0.522], *p_adj_*= < 0.001; 0 vs 1% 12-15m, Cliff’s delta = 0.728, 95CI [0.749, 0.707], *p_adj_* = < 0.001). Shoal kinesis changed negligibly over time with 0.5% alcohol treatment, while it was reduced almost immediately with 1% alcohol treatment. The effect became larger over time (Figure 4E, 0 vs 1%, 6-9m, Cliff’s delta = - 0.419, 95CI [−0.387, −0.449], *p_adj_* = 0.001). Thus, immersion in 0.5% alcohol reduced anxiety-like behaviours, while sedative-like effects were more evident when immersed in 1% alcohol (Figure 4D, 4E).

The *chrna3* mutants seemed to experience the effects of alcohol exposure differently. Behaviour of unexposed (0% alcohol) was similar between the two genotypes with the *chrna3* exhibiting only slight differences in shoal cohesion (Figure S7, A), and shoal kinesis (Figure S7B). However, unlike the WT shoals, exposure to 0.5% alcohol reduced shoal cohesion only marginally (Figure 4F) and it did not increase over time as it did in the WTs (Figure 4D). This difference is clearer when genotypes are compared directly. Mutant fish retained significantly tighter shoaling cohesion compared to the wild type (Figure 4H, 12-15m, Cliff’s delta = −0.44, 95CI [−0.409, −0.47], *p_adj_* = < 0.001). Meanwhile, shoal kinesis was initially heightened in *chrna3* fish compared to the WTs (Figure 4I, 0-3m, Cliff’s delta = 0.253, 95CI [0.288, 0.219], *p_adj_*= 0.004), which paralleled the *chrna3* alcohol-induced motility stimulation in the SAZA (Figure S5, A-F). The alcohol-induced kinesis difference between genotypes gradually declined as the assay progressed (Figure 4I). This divergence in behaviours between genotypes suggests that the anxiolysis from the alcohol exposure was much lower in *chrna3* than that experienced by their WT counterparts.

The behaviour differences between the mutant and wild-type shoals were minimal in 1% alcohol for the majority of the assay duration (Figure S7, C-D). However, in the latter part of the assay mutants seemed to resist the sedative effects of alcohol, retaining higher shoal kinesis than the wild type (Figure S7D, 15-18m, Cliff’s delta = 0.372, 95CI [0.405, 0.34], *p_adj_* = 0.008). The effects of alcohol on shoal cohesion and kinesis in the first five minutes of exposure were similar between genotypes (Figure 4, S7). Overall, these results from the shoaling assay suggest that *chrna3* had reduced effect of alcohol exposure on anxiolysis as well as sedative effects seen in WT fish.

### Whole brain transcriptomic analysis shows major neurotransmitter signalling pathways are affected in mutants

To gain an understanding of these behavioural differences we performed bulk RNA sequencing of WT and mutant whole brains. Differential gene expression (DEG) analysis suggested that the two genotypes had large differences, with PC1 accounting for ∼44% of the variation in a PCA (Figure 5A). Gene Set Enrichment Analysis (GSEA) revealed notable enrichment in transcriptional changes in mutants (Figure 5B). The significantly enriched biological functions impacted in the mutant included neuronal development, ion channel function, and secondary messenger-associated processes (Figure 5B). Additionally, genes associated with feeding behaviour were also enriched. In total, the expression of ∼1600 genes was significantly changed (2-fold or more), and a further ∼4600 genes were identified with provisional alterations (less than 2 fold; Figure 5C). Of these, there was an approximately even split between up- and down-regulated gene numbers (Figure 5C). Together, this perhaps indicates a wide cascade of compensatory changes induced by the reduced expression and function of *chrna3*.

**Figure 5.**
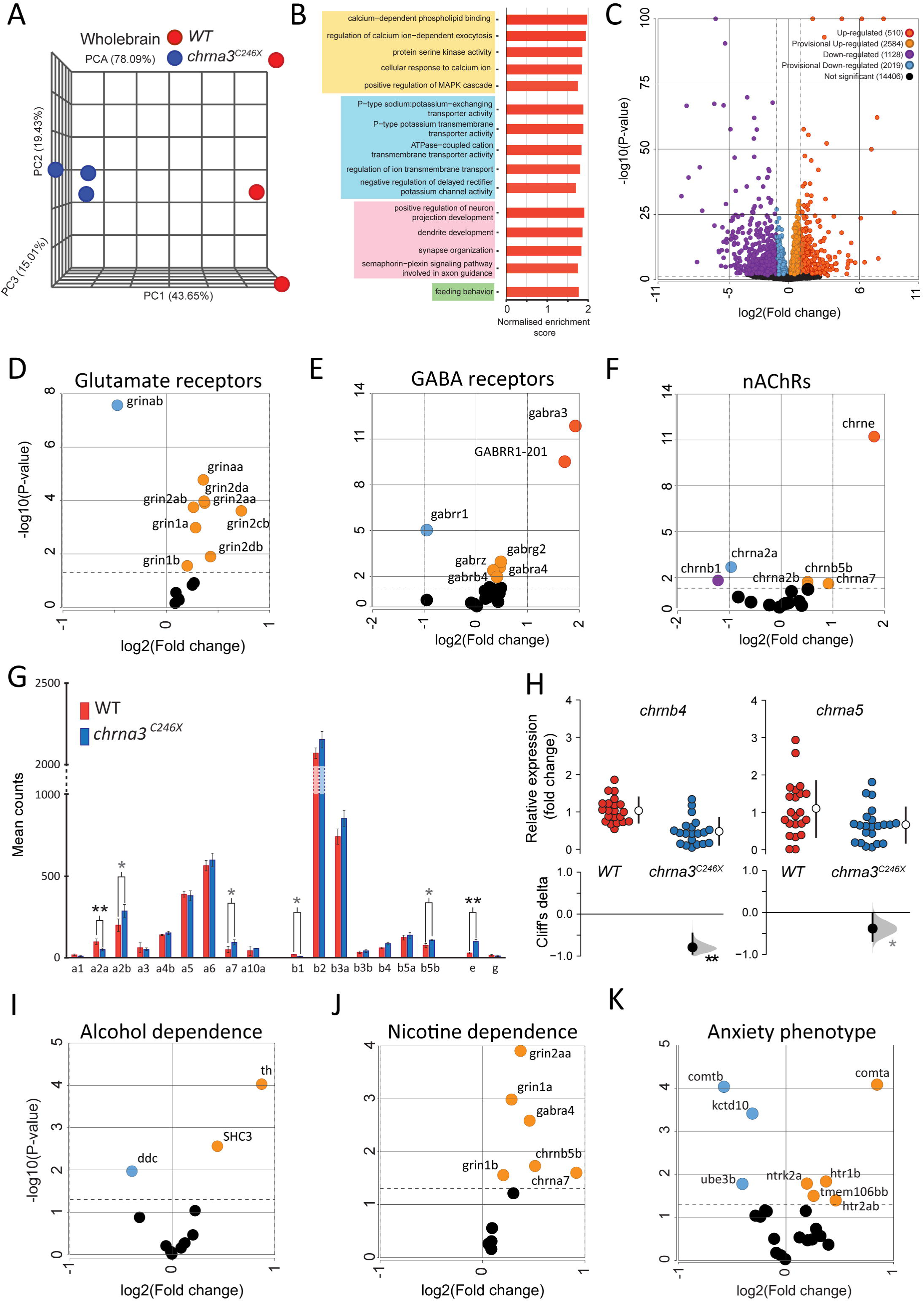
*chrna3^C246C^* mutants exhibit transcriptional changes in dependence, neuronal development, and ion channel process associated genes. RNAseq comparison of *chrna3^C246C^* (n = 3) vs *chrna3^C246C^* (n = 3) 12 week old zebrafish whole brain tissues. ***A.*** PCA clustering of samples by genotype. ***B.*** All significantly enriched GO terms between genotypes. Positive enrichment values indicate enrichment in the *chrna3^C246C^* mutant zebrafish. Significance thresholds are defined at *p* = < 0.05 and FDR < 0.2. Yellow = secondary messenger associated, Blue = ion channel associated, pink = neuronal differentiation and development, green = behaviours. ***C.*** Volcano plot of gene expression changes between genotypes. Up/down-regulated genes = *p* < 0.05 & log2 fold change > ±2. Provisional up/down-regulated genes = *p* < 0.05 & log2 fold change < ±2. Not significant genes = *p* > 0.05. The colour key applies to D-F and I-K. Positive fold change values indicate up-regulation in the *chrna3^C246C^* mutant zebrafish. ***D-F.*** Subsets of volcano plot ***C***, highlighting ***D.*** glutamate, ***E.*** GABA, and ***F.*** nicotinic acetylcholine (nAChR) receptor gene families. ***G.*** Mean counts (± standard deviation) of nAChRs (gene prefix ‘chrn’). Stars indicate significant difference between genotypes, ** = p < 0.01, * = p < 0.05. ***H.*** Fold change in WT vs *chrna3^C246C^* of *chrna5* (n = 21/20) and *chrnb4* (n = 22/22) by qRT-PCR. Stars indicate significant difference between genotypes, ** = p < 0.01, * = p < 0.05. ***I-K.*** Subsets of volcano plot ***C***, highlighting genes associated with ***I.*** nicotine or ***J.*** alcohol dependence (Hu et al., 2018), and ***K.*** anxiety phenotypes (Howe et al., 2016; Koskinen and Hovatta, 2023). The full gene lists can be found in Table 4.

Major neurotransmitter signalling including glutamate, and GABA receptors in addition to nAChRs, which are activated by exposure to psychoactive substances like alcohol, also showed modest changes in expression patterns (Figure 5D-F). Both glutamate and GABA receptors appeared to show a small but consistent upregulation amongst many receptor subunits (Figure 5D-E). The expression of most nAChR genes however was unchanged, though a few were positively or negatively regulated with no consistent trend (Figure 5F). Closer inspection of the nAChR subunit gene counts revealed that expression was very low for some subunits, as was highlighted for *chrna3* in the HCR (Figure 1, 5G). For genes with low expression counts it is a challenge to determine significant changes through RNA-seq analysis. As *CHRNA3* expression is co-regulated with *CHRNA5* and *CHRNB4* in humans, both of which have also been associated with substance use disorders (Wen et al., 2016), we examined whether the expression of these genes had also been altered in the mutant. Both *chrna5* and *chrnb4* displayed significant downregulation by qPCR, with the effect size being much larger for the latter (Figure 5H).

We also examined gene sets curated from meta-analyses of genes consistently associated with alcohol and nicotine dependence in humans (Hu et al., 2018), and with mammalian anxiety phenotypes (Howe et al., 2016; Koskinen and Hovatta, 2023) to address whether *chrna3* dysfunction also alters expression of these genes (Figure 5I-K). The alcohol dependence-associated genes exhibited few changes (Figure 5I), however, ∼50% of the nicotine-dependence genes were provisionally upregulated in the mutant (Figure 6J). Finally, genes associated with anxiety phenotypes also showed a number of expression changes, albeit some in positive and some in negative direction of expression (Figure 5K). In all cases, this analysis revealed that though the expression of a large number of these genes changed they were modest.

## Discussion

In this study, a refinement of self-administration zebrafish assay (SAZA) revealed an acute biphasic response to alcohol, in juvenile zebrafish, with preference transitioning to avoidance when exposure increased to ∼0.7% alcohol (Figure 2). This biphasic response to alcohol aligns zebrafish with other animals, suggesting that hormesis to alcohol is a fundamental property across vertebrates (Figures 1, 2; (Calabrese and Baldwin, 2003; Calabrese, 2008; Sterling et al., 2015)). The closed-loop design of SAZA, similar to a two-bottle choice assay used for rodents, facilitated the analysis of uninhibited choice and isolated *chrna3*’s role in modulating this biphasic response.

Isolating contributions of individual genetic factors to a phenomena such as substance dependence is challenging. It is further compounded for multi-subunit receptor genes that cluster at a genomic locus, such as the *CHRNA5-A3-B4* gene cluster. This genomic locus has been consistently associated with several substance use disorder GWASes (Chen et al., 2009; Maes et al., 2011; Haller et al., 2014; Wen et al., 2016). Although, male transgenic mice overexpressing *Chrna5-a3-b4* consume less alcohol than wild-type mice (Gallego et al., 2012), a causal relationship can’t be ascribed to *Chrna3* alone from this study (Kamens et al., 2023). *Chrna3* knockout mice die prematurely as neonates (Xu et al., 1999) and the conditional knockout (Elayouby et al., 2021) only examined nicotine response leaving the role of α3 in alcohol stimulation unclear (Kamens et al., 2009). Zebrafish *chrna3* mutants, however, survive to adulthood, offering a tractable system to investigate gene-specific effects here. *chrna3* mutants displayed a blunted aversive switch in the biphasic response, while maintaining early attraction (Figure 2, 3). WT fish exhibited aversive behaviour to 0.7% or higher alcohol stimulus, which only became observable in mutants at higher concentrations. This aligns with rodent studies, where a broad spectrum nicotinic receptor antagonist 18-MC in the medial habenula (MHb) - interpeduncular nucleus (IPN) circuit reduced transmission and increased nicotine or alcohol self-administration, respectively (Glick et al., 2011; Miller et al., 2019). As this circuit co-expresses all the three nAChR subunits α3, α5, and β4, they are thus proposed to mediate aversion to psychoactive substances, including alcohol (Marks et al., 1992; Xu et al., 2006; Frahm et al., 2011; Fowler and Kenny, 2012; Mathuru, 2018; Elayouby et al., 2021). However, as shown in the gene expression profile here (Figure 1), and in a recent report (Hua et al., 2025), *chrna3* is expressed widely in the central and peripheral nervous system. Consequently, altered alcohol tolerance reported here may stem from a combined effect across multiple circuits and systems. Further to this limitation, although most physiological features and brain regions described in adult zebrafish are present in juveniles by 25 dpf, late-stage developmental changes may influence adult phenotypes (Schmidt et al., 2013; Singleman and Holtzman, 2014; Raj et al., 2020), warranting further studies.

A shifted tolerance window exhibited by *chrna3* mutants (Figure 3C) suggests a threshold based engagement of different brain circuits, similar to the proposed mammalian twin threshold model (Grasing, 2016). This model posits that cholinergic signalling, particularly in the VTA and nucleus accumbens (NAc), modulates drug reinforcement behaviours. In this model, lower threshold increases positive reward association probability, stimulating behavioural repetition, while a second, higher threshold does the opposite as different circuits come into play (Grasing, 2016). This may also be applicable to other cholinergic brain regions, such as the MHb. Additionally, higher agonist requirements to activate α5, α3, and β4 containing nAChRs (Coe et al., 2005), could elevate the cholinergic threshold to engage this aversive circuit, providing a potential explanation for why *chrna3* mutation predominantly affected the avoidance response.

Beyond preference, *chrna3* mutation also blunted the psychoactive effects of alcohol during immersion. Immersion is a rapid, non-invasive route for administration of water-soluble pharmaceutics, uniquely available for aquatic animals. Alcohol immersion time correlates with blood alcohol concentration (BAC) until equilibrium is reached, and is a key measure affecting physiological responses (Lockwood et al., 2004). As in humans, BAC > 0.05%, or the equivalent of ∼2-3 Australian “standard drinks”, results in significant behavioural impairment in zebrafish, which can be reached within 1-2 minutes of immersion in 1.5% alcohol. This manifests in humans as slower reaction times (Grant et al., 2000), and zebrafish as loss of swimming coordination or impaired locomotion (Tran and Gerlai, 2013; Tsang et al., 2019). However, lower concentrations are stimulatory, activating dopaminergic reward pathways and increasing locomotion, while reducing anxiety-like behaviours and aggression (Echevarria et al., 2010; Araujo-Silva et al., 2020).

Shoaling behaviour, that becomes prominent by 14 dpf (Miller and Gerlai, 2007), can effectively model these contrasting effects (Oscar-Berman and Marinković, 2007; Kurta and Palestis, 2010; Miller et al., 2013; Rueger and King, 2013) at SAZA-relevant concentrations. In wild types, both 0.5% and 1% alcohol immersion gradually reduced shoal cohesion, but exhibited differing effects on shoal kinesis (Figure 4). One interpretation of the coupled reduction in cohesion and kinesis in 1% alcohol is as a sedative effect, evident also in the video recordings (Movie 2). Meanwhile, reduction of cohesion alone could indicate an anxiolytic effect, that dominated in a narrow alcohol concentration range (0.5%). These concentrations approximately correlated with the attractive and aversive doses during SAZA, suggesting a potential link. However, the *chrna3* mutants displayed no reduction in shoal cohesion at 0.5% (Figure 4H). Whether this reduction in anxiolysis phenotype in the mutants was directly due to reduced sensitivity to alcohol or independent of it needs further investigation. Irrespective, it is evident that the *chrna3* mutation blunted drug-induced behavioural effects, including the reduced aversion in SAZA.

Alcohol-induced kinesis also differed between genotypes, with *chrna3* mutants exhibiting stronger alcohol-induced locomotion in SAZA and shoaling assays (Figure 4, S5). This may be correlated to the reduced expression of *chrna3* in the caudal and rostral regions of the mutant facial motor nucleus (Fmn), also termed the octavolateral efferent nucleus (OEN) (Figure 1, S1), a hindbrain region connected to the lateral line. Cholinergic signalling here is hypothesised to act as a ‘brake’ on locomotor auto-sensory stimulation generated during swimming (Pichler and Lagnado, 2020; Manuel et al., 2021; Odstrcil et al., 2022). While typically this function is assigned to α9 receptors, if α3 plays a role, as suggested recently (Rima et al., 2020), it may account for the heightened alcohol-induced locomotor response in *chrna3* mutants. However, *chrna3*’s influence on locomotor control requires further investigation, as rodent studies offer inconsistent evidence (Kamens et al., 2009).

Whole-body mutation in our *chrna3* mutant precluded implicating specific brain regions like the dorsal habenula (MHb in mammals), yet, it does inform on potential consequences of *CHRNA3* variants that reduce α3 function in humans. Whole brain RNA sequencing identified broad transcriptional profile change in the mutants. Small, but, consistent upregulation of nicotine dependence associated genes was indicative of a bi-directional relationship (Figure 5J; (Hu et al., 2018)), aligning with GWAS studies linking *CHRNA3* polymorphisms to increased nicotine dependence (Chen et al., 2009; Maes et al., 2011). Downregulation of *chrna5* and *chrnb4* (Figure 5H), also associated with substance dependence (Fowler et al., 2011; Frahm et al., 2011) and co-expressed with *chrna3* in other vertebrates, potentially contributed to the mutant’s alcohol tolerance. Generation and characterization of an independent *chrna3*, *chrna5,* and *chrnb4* mutant lines along with their combinations (currently in progress), will be invaluable to dissociate these factors further.

Beyond nAChRs, alcohol impacts both glutamate and GABA receptors, modulating excitatory and inhibitory tones, respectively (Davies, 2003; Möykkynen and Korpi, 2012; Jones, 2019). Consequently, changes in expression of these receptor genes (Figure 5D, 5E) in mutants was also interesting to observe, as many homologs also feature in nicotine dependence and anxiety phenotypes in humans (Figure 5J, 5K). At a functional level, neuronal receptor related GO terms like synaptic signalling, ion channel function, and cation homeostasis were significantly enriched (Figure 5B). However, neural development terms including neuron projection, dendrite development, and axon guidance were also enriched despite the use of adult brains (Figure 5B), perhaps indicative of compensatory plasticity. Finally, enrichment of feeding behaviour associated genes was notable (Figure 5B), as cholinergic signalling and CHRNA5-CHRNA3-CHRNB4 cluster SNPs have been tentatively linked by GWAS studies to smoking/obesity interactions (Varga et al., 2013; Herman et al., 2016; Ortiz-Guzman et al., 2024). This transcriptomic change warrants further examination to determine the simultaneous influence of *chrna3* on substance dependence and other comorbid disorders.

Overall, these findings suggest the type of brain-wide compensatory changes that may occur corresponding to loss of *chrna3* function, eventually resulting in heightened alcohol tolerance and potential predisposition to multi-substance use and comorbid disorders.

Understanding alcohol’s biphasic properties is imperative to address Alcohol Use Disorder (AUD). Our study fills a gap, providing novel evidence on the role of *chrna3* in modulating alcohol preference, and tethering it to a specific facet of the dose-response curve. Given the paucity of animal model data, despite GWAS associations (Haller et al., 2014), the evidence of isolated gene function from a whole body mutation in zebrafish presented here represents a valuable step forward in the context of phenotypes leading to alcohol dependence development in humans harbouring *CHRNA3* variants.

## Acknowledgements

We would like to thank the zebrafish fish facility (ZFF) staff at the IMCB, A*STAR for their assistance with fish husbandry. We also thank Li Xinrui for generating customized Python analysis scripts. Additionally, during the preparation of this work the author(s) used Grammarly, and Gemini in order to assist with grammar and consistency of phrasing. After using this tool/service, the author(s) reviewed and edited the content as needed and take(s) full responsibility for the content of the publication.

## Supplementary Materials

Supplementary Material Figures S1 to S7, Tables S1 to S3. Number of supplementary figures and tables = 10

## Author Contributions

Conceptualization, A.S.M, and J.R; methodology, A.S.M., T.D.B., C.K., and J.R.; formal analysis, J.R., T.D.B and A.S.M.; investigation, J.R., C.K., and A.S.M; resources, A.S.M; writing—original draft preparation, J.R, and A.S.M.; writing—review and editing, All.; visualisation, J.R.; supervision, A.S.M.; project administration, A.S.M., A.M.; funding acquisition, A.S.M.

## Funding

This research was funded by the Ministry of Education (MOE), Singapore (through grant number T2EP30220-0020), and Yale-NUS College (through grant numbers IG16-LR003, IG18-SG103, IG19-BG106, and SUG) to ASM.

## Data Availability Statement

The datasets used to generate the figures are included within the article and its additional file(s). Additional data are available from the corresponding author upon request.

## Conflicts of Interest

The authors declare no conflict of interest. The sponsors had no role in the design, execution, interpretation, or writing of the study.

## Figure & Table Legends

**Table 1.** Primers for single-guide RNA template. CRISPR targets are underlined. Homology arms in lower case.

**Table 2.** Mean time spent by zebrafish in the stimulus chamber during SAZA over an 18 minute stimulus dispensing period (n = 30-32)

**Table 3.** Primers used for qPCR of *chrna3*, *chrna5*, *chrnb4*, and housekeeping genes.

**Table 4.** Probe sequences for FISH-HCR. B1 Probes were visualised with Alexa fluorophore 546, B2 probes were visualised with Alexa fluorophore 647, and B3 probes were visualised with Alexa fluorophore 488. P1 and P2 probes of the same gene are paired according to number 1-8.

**Table 5.** Genes included in the custom sets ‘Nicotine dependence’ and ‘Alcohol dependence’. These genes have been associated with nicotine and alcohol dependence, described by Hu *et al*. <u>(Hu et al. 2018)</u>

## Movie Legends

**Movie 1:** A representatve segment of the zebrafish self-administraton assay (SAZA) with 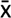1.5% alcohol shown in Figure 1B.

**Movie 2:** A representative segment of the shoaling assay with 1% alcohol shown in figure 4D.

## Supplementary Figure & Table Legends

**Figure S1.** Replicates of *chrna3^C246C^* expression from figure 1, in *WT* (*A*) and *chrna3^C246C^* (***B***) 14dpf zebrafish larvae facial motor nucleus (Fmn) marked by *chata* expression, visualised by *in-situ* hybridisation chain reaction (HCR). Scale bars = 10 μm. ***C.*** Comparison of the mean fluorescent intensity (± standard deviation) of *chrna3^C246C^* in the demarcated zone between genotypes (Mean difference = −24.0 [95CI −26.3,- 20.9], p = 0.012).

**Figure S2.** Replicates of *chrna3^C246C^* expression from figure 1, in *WT* (***A***) and *chrna3^C246C^* (***B***) 14dpf zebrafish larvae right dorsal habenula (Hb) marked by *nrp1a* expression, visualised by *in-situ* hybridisation chain reaction (HCR). Scale bars = 20 μm. ***C.*** Comparison of the mean fluorescent intensity (± standard deviation) of *chrna3^C246C^* in the demarcated zone between genotypes (Mean difference = 0.168 [95CI −2.84, 1.73], p = 0.944).

**Figure S3.** Gardener-Altman and Cumming estimation plots show alcohol response in three-minute intervals. No stimulus was dispensed in the ‘Pre’ and ‘Post’ periods. Time spent in the stimulus zone during 0% SAZA condition for ***A.*** WT (n = 31) and ***B.*** *chrna3^C246C^* mutants (n = 32). Time spent in the control zone for ***C, E, G.*** WT (n = 30/31 /30) and ***D, F, H****. chrna3^C246C^* mutants (n = 29/32/30) in ***C, D.*** 0%, ***E, F.*** 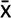0.5% and ***G, H.*** 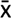1.5% SAZA. Asterisks indicate a significant difference between that time portion and the pre-stimulus period: * = p_adj_ < 0.05 ** = p_adj_ < 0.01. See Table S1 for the precise effect sizes and p-values, corrected for multiple comparisons.

**Figure S4.** Gardener-Altman and Cumming estimation plots show alcohol response in three-minute intervals. No stimulus was dispensed in the ‘Pre’ and ‘Post’ periods. Number of entries into the stimulus zone for ***A-C.*** WT (n = 30/31 /30) and *D-F. chrna3^C246C^* mutants (n = 29/32/30) in ***A, D.*** 0%, ***B, E.*** 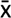0.5% and ***C, F.*** 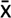1.5% SAZA. Asterisks indicate a significant difference between that time portion and the pre-stimulus period: * = p_adj_ < 0.05 ** = p_adj_ < 0.01. See Table S1 for the precise effect sizes and p-values, corrected for multiple comparisons.

**Table S1**. Statistical analysis output table for Figure 2 A-E, and Figures S3 and S4, intragenotype paired mean difference of % time in stimulus, time in control, or number of entries, for repeated measures against the ‘Pre’ period for 0%, x0.5% and 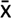1.5% alcohol SAZA.

**Figure S5.** Gardener-Altman and Cumming estimation plots for total SAZA self-administration period for WT (***A, D***), *chrna3^C246C^* (***B, E***) and comparisons (***C, F***). Plots display the comparison of velocity inside (***A-C***) or outside (***D-F***) the stmulus zone at different concentratons (0/x0.5/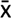1.5%) for WT (n = 30/31/30) and *chrna3^C246C^* zebrafish (n = 29/32/30). ***G-I.*** Gardener-Altman and Cumming estimation plots for ***G.*** 0%, ***H.*** 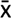0.5%, and ***I.*** 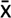1.5% SAZA. Plots compare preference index (PI) between WT (0/x0.5/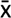1.5% n = 30/ 31/30) and *chrna3^C246C^* (0/x0.5/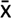1.5% n = 29/32/30) within each time period. No stimulus was dispensed in the ‘Pre’ and ‘Post’ periods. Asterisks indicate the following significant differences from 0% alcohol treatments (***A, B, D, E***) or WT (***C, F-I***): * = p < 0.05, and effect size reported by Cliff’s delta between ± 0.2 and ± 0.4 are considered provisional, ** = p < 0.01 and effect size bigger than ± 0.4 is considered a meaningful difference. See Table S2 for exact effect size and p-values.

**Figure S6**. Cliff’s delta (± 95% CI) forest plots of all calculated metrics from 0/x0.5/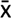1.5% alcohol SAZA, comparing the response of wild type to *chrna3^C246C^* mutant zebrafish. Positive values indicate a greater response in the mutant. See Table S2 for exact effect size and p-values.

**Table S2.** Statistical analysis output table for figure 3A, B, intragenotype, and figure 3C, intergenotype, comparisons of stimulus volume dispensed over the full self administration period for 0, x0.5 and 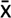1.5% alcohol SAZA. Statistical analysis output table for figure 3 D-I, paired mean difference and delta-delta in preference index (PI) between time periods, Figure S5 A-C, comparing preference index (PI), velocity, and Figure S6 other metrics for 0, x0.5 and 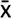1.5% alcohol SAZA.

**Figure S7**. *WT and chrna3^C246C^ mutants share shoaling cohesion and kinesis phenotypes during 0% and 1% alcohol immersion.* Mean (± 95% CI) shoal cohesion (***A, C***) and kinesis (***B, D***) with Cumming estimation plots comparing 0% (***A, B***), or 1% alcohol exposure (***C, D***) between genotypes. Metrics compared per frame at 2fps (n = 6 assays per genotype, per treatment). Asterisks indicate the following significant differences from WT within that time period: * = p < 0.05, and effect size reported by Cliff’s delta between ± 0.2 and ± 0.4 are considered provisional, ** = p < 0.01 and effect size bigger than ± 0.4 is considered a meaningful difference. See Table S3 for exact effect size and p-values.

**Table S3.** Statistical analysis output table for figure 4A-G, comparing shoaling behaviours within genotype, figure 4H-I and S7, between genotypes, displaying pairwise effect size comparisons, and repeated measures estimated marginal mean difference adjusted p values within time period.

## Notes

### Competing Interest Statement

The authors have declared no competing interest.

### Summary of Updates

Updated figures with matching revised supplementary figures. Introduction, results, and discussion sections updated for clarity.

