## Supplementary Material for "*chrna3* modulates alcohol response"

**Figure S1 to S10**

**Tables S1 to S16**

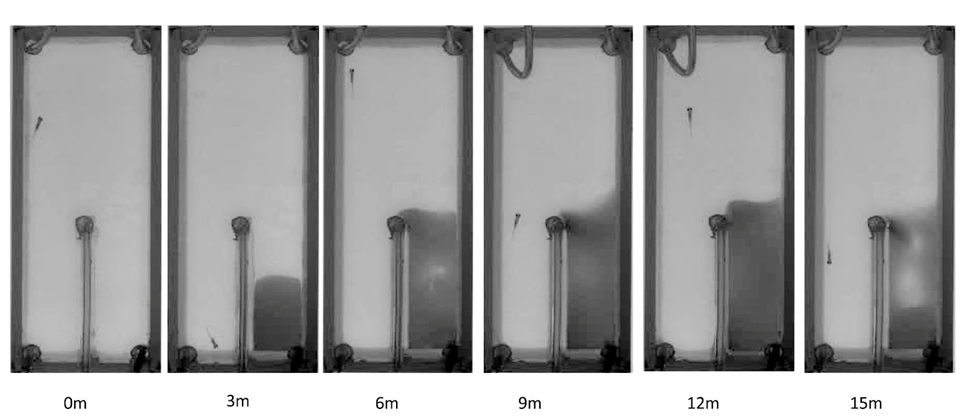
**Figure S1. The SAZA system maintains chemically distinct but physically connected zones.** Demonstration SAZA using food dye in as the stimulus dispensing solution to indicate the creation and restriction of the chemically distinct stimulus zone.

Table S1. Mean time spent by zebrafish in the stimulus chamber during SAZA over an 18 minute stimulus dispensing period (n = 30-32)

| **Solution dispensed** | **Mean time in stimulus chamber (seconds)** |
| --- | --- |
| 0% alcohol | 238 |
| 20% alcohol (x̄0.5%) | 220 |
| 70% alcohol (x̄1.5%) | 132 |

Table S2. Statistical analysis output table for figure 2 A-E

| **Genotype & alcohol treatment (Figure)** | **Time period vs PRE (mins)** | **Cliff's Delta** | **95% CI upper** | **95% CI lower** | **P value** |
| --- | --- | --- | --- | --- | --- |
| WT x̄0.5% (3A) | 0-3 | 0.107 | 0.388 | -0.197 | 0.473 |
| WT x̄0.5% (3A) | 3-6 | 0.403 | 0.636 | 0.093 | 0.007 |
| WT x̄0.5% (3A) | 6-9 | 0.211 | 0.478 | -0.080 | 0.152 |
| WT x̄0.5% (3A) | 9-12 | -0.159 | 0.147 | -0.446 | 0.285 |
| WT x̄0.5% (3A) | 12-15 | -0.376 | -0.082 | -0.617 | 0.009 |
| WT x̄0.5% (3A) | 15-18 | -0.365 | -0.082 | -0.617 | 0.009 |
| WT x̄0.5% (3A) | POST | -0.38398 | -0.07388 | -0.62539 | 0.007 |
| WT x̄1.5% (3B) | 0-3 | 0.0844 | 0.376 | -0.224 | 0.571 |
| WT x̄1.5% (3B) | 3-6 | 0.0511 | 0.349 | -0.24 | 0.724 |
| WT x̄1.5% (3B) | 6-9 | -0.424 | -0.12 | -0.651 | 0.005 |
| WT x̄1.5% (3B) | 9-12 | -0.658 | -0.404 | -0.822 | < 0.001 |
| WT x̄1.5% (3B) | 12-15 | -0.471 | -0.178 | -0.696 | 0.002 |
| WT x̄1.5% (3B) | 15-18 | -0.653 | -0.389 | -0.829 | < 0.001 |
| WT x̄1.5% (3B) | POST | -0.24 | 0.0867 | -0.518 | 0.12 |
| C246X x̄0.5% (3D) | 0-3 | 0.186 | 0.465 | -0.109 | 0.193 |
| C246X x̄0.5% (3D) | 3-6 | 0.524 | 0.740 | 0.233 | < 0.001 |
| C246X x̄0.5% (3D) | 6-9 | 0.568 | 0.762 | 0.281 | < 0.001 |
| C246X x̄0.5% (3D) | 9-12 | 0.334 | 0.588 | 0.039 | 0.018 |
| C246X x̄0.5% (3D) | 12-15 | 0.213 | 0.490 | -0.088 | 0.127 |
| C246X x̄0.5% (3D) | 15-18 | 0.273 | 0.535 | -0.027 | 0.056 |
| C246X x̄0.5% (3D) | POST | 0.048828 | 0.347656 | -0.24219 | 0.7294 |
| C246X x̄1.5% (3E) | 0-3 | 0.157778 | 0.44 | -0.15778 | 0.2952 |
| C246X x̄1.5% (3E) | 3-6 | 0.117778 | 0.41111 | -0.1866 | 0.4282 |
| C246X x̄1.5% (3E) | 6-9 | -0.22889 | 0.082222 | -0.49333 | 0.1276 |
| C246X x̄1.5% (3E) | 9-12 | -0.39111 | -0.1 | -0.64 | 0.0082 |
| C246X x̄1.5% (3E) | 12-15 | -0.44667 | -0.15556 | -0.68 | 0.0032 |
| C246X x̄1.5% (3E) | 15-18 | -0.3622 | -0.05778 | -0.61556 | 0.0136 |
| C246X x̄1.5% (3E) | POST | -0.33778 | -0.02667 | -0.58889 | 0.0252 |

Table S3 Statistical analysis output table for figure 2F

| **Metric (Figure)** | **Alcohol treatment (vs WT)** | **Cliff's Delta** | **95% CI upper** | **95% CI lower** | **P value** |
| --- | --- | --- | --- | --- | --- |
| Volume dispensed (3F) | x̄0.5% | 0.328 | 0.584 | 0.009 | 0.027 |
| Volume dispensed (3F) | x̄1.5% | 0.188 | 0.466 | -0.123 | 0.206 |

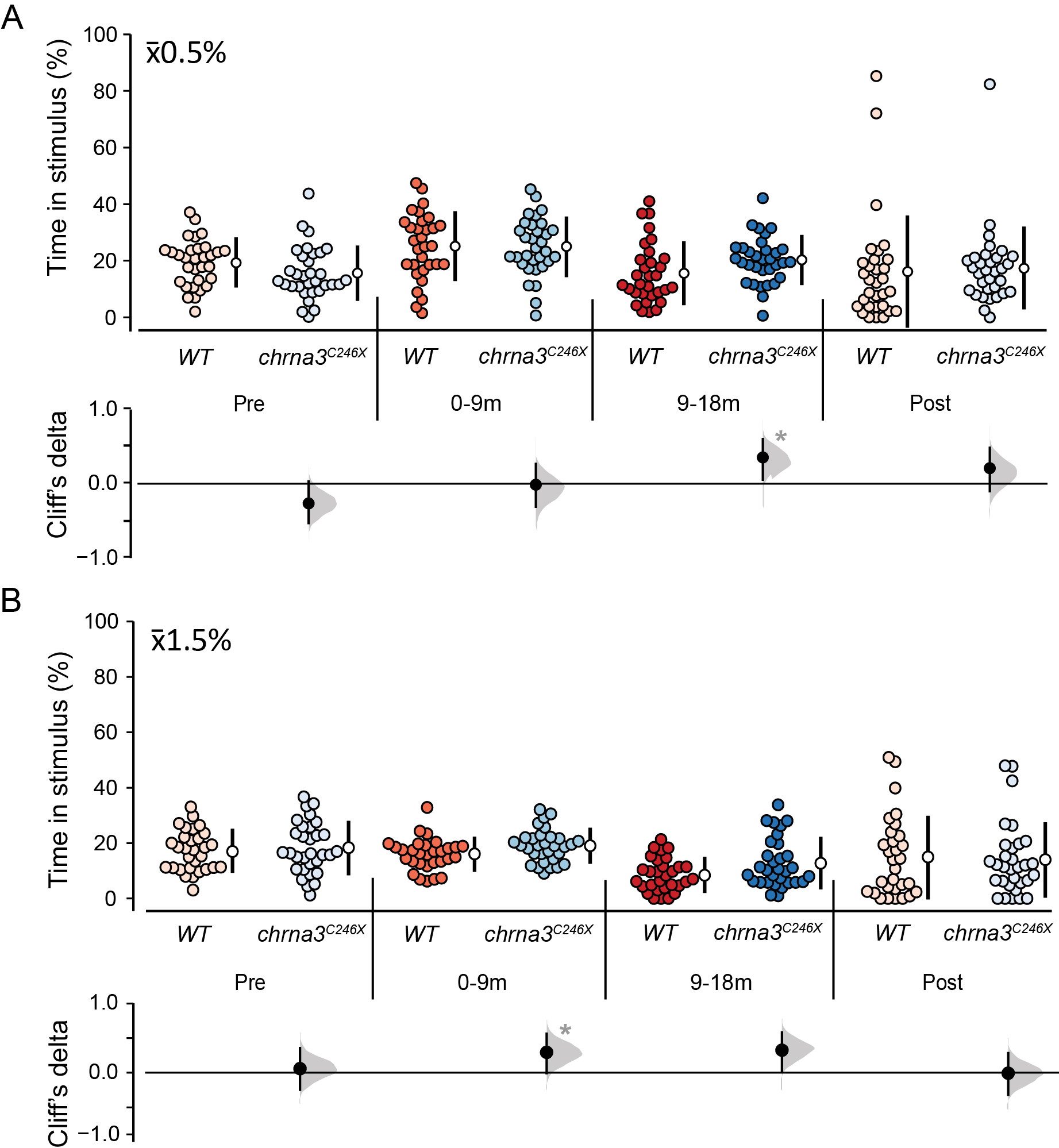

**Figure S2. *chrna3^C246X^*** **mutants exhibit an increased alcohol tolerance window.** Gardener-Altman and Cumming estimation plots for x̄0.5% (**A**) and x̄1.5% (**B**) SAZA. Plots compare time spent in the stimulus zone between WT (x̄0.5/x̄1.5% n = 31/30) and *chrna3^C246X^* (x̄0.5/x̄1.5% n = 31/30) within each time period. No stimulus was dispensed in the ‘Pre’ and ‘Post’ periods. Asterisks indicate the following significant difference between genotypes: * = *p* < 0.05, and effect size reported by Cliff’s delta between ± 0.2 and ± 0.4 are considered provisional, ** = p < 0.01 and effect size bigger than ± 0.4 is considered a meaningful difference. See table S4 for exact effect size and p-values.

Table S4 Statistical analysis output table for figure S2

| **Alcohol treatment (Figure)** | **Time point (vs WT)** | **Cliff's Delta** | **95% CI upper** | **95% CI lower** | **P value** |
| --- | --- | --- | --- | --- | --- |
| x̄0.5% (S2A) | Pre | -0.26613 | 0.030242 | -0.52823 | 0.065 |
| x̄0.5% (S2A) | Stim start 0-9 | -0.01613 | 0.268145 | -0.30847 | 0.910 |
| x̄0.5% (S2A) | Stim end 9-18 | 0.352823 | 0.602823 | 0.054435 | 0.015 |
| x̄0.5% (S2A) | Post | 0.207661 | 0.481855 | -0.10081 | 0.164 |
| x̄1.5% (S2B) | Pre | 0.058065 | 0.354839 | -0.24516 | 0.691 |
| x̄1.5% (S2B) | Stim start 0-9 | 0.296774 | 0.56129 | -0.0086 | 0.047 |
| x̄1.5% (S2B) | Stim end 9-18 | 0.270968 | 0.522581 | -0.04301 | 0.066 |
| x̄1.5% (S2B) | Post | -0.0129 | 0.283871 | -0.32043 | 0.925 |

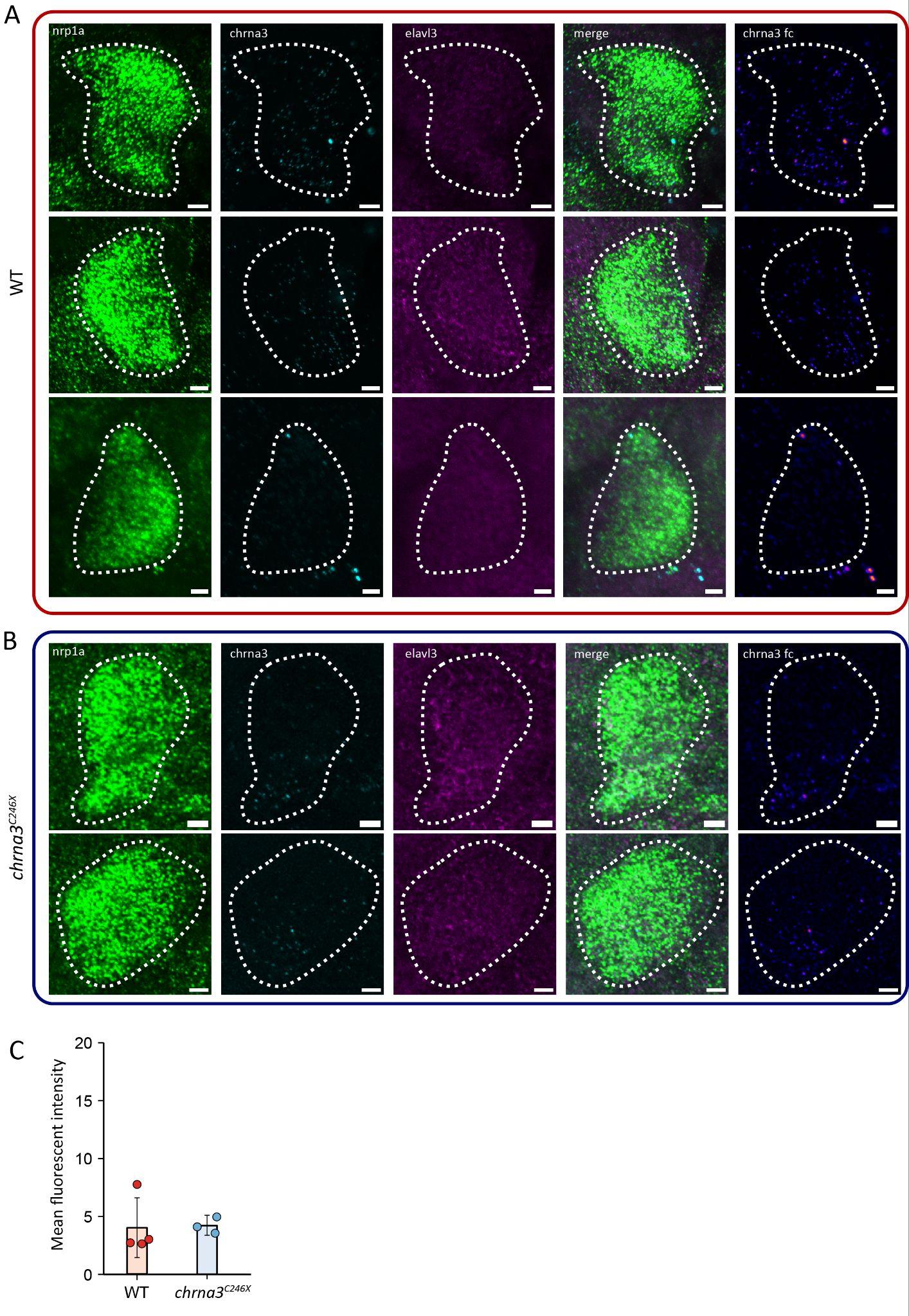

**Figure S3.** *chrna3* expression in *WT* (**A**) and *chrna3^C246X^* (**B**) 14dpf zebrafish larvae right dorsal habenula (Hb) marked by *nrp1a* expression, visualised by *in-situ* hybridisation chain reaction (HCR). **C.** Comparison of the mean fluorescent intensity (± standard deviation) of *chrna3* in the demarcated zone between genotypes (Mean difference = 0.168 [95CI -2.84, 1.73], p = 0.944).

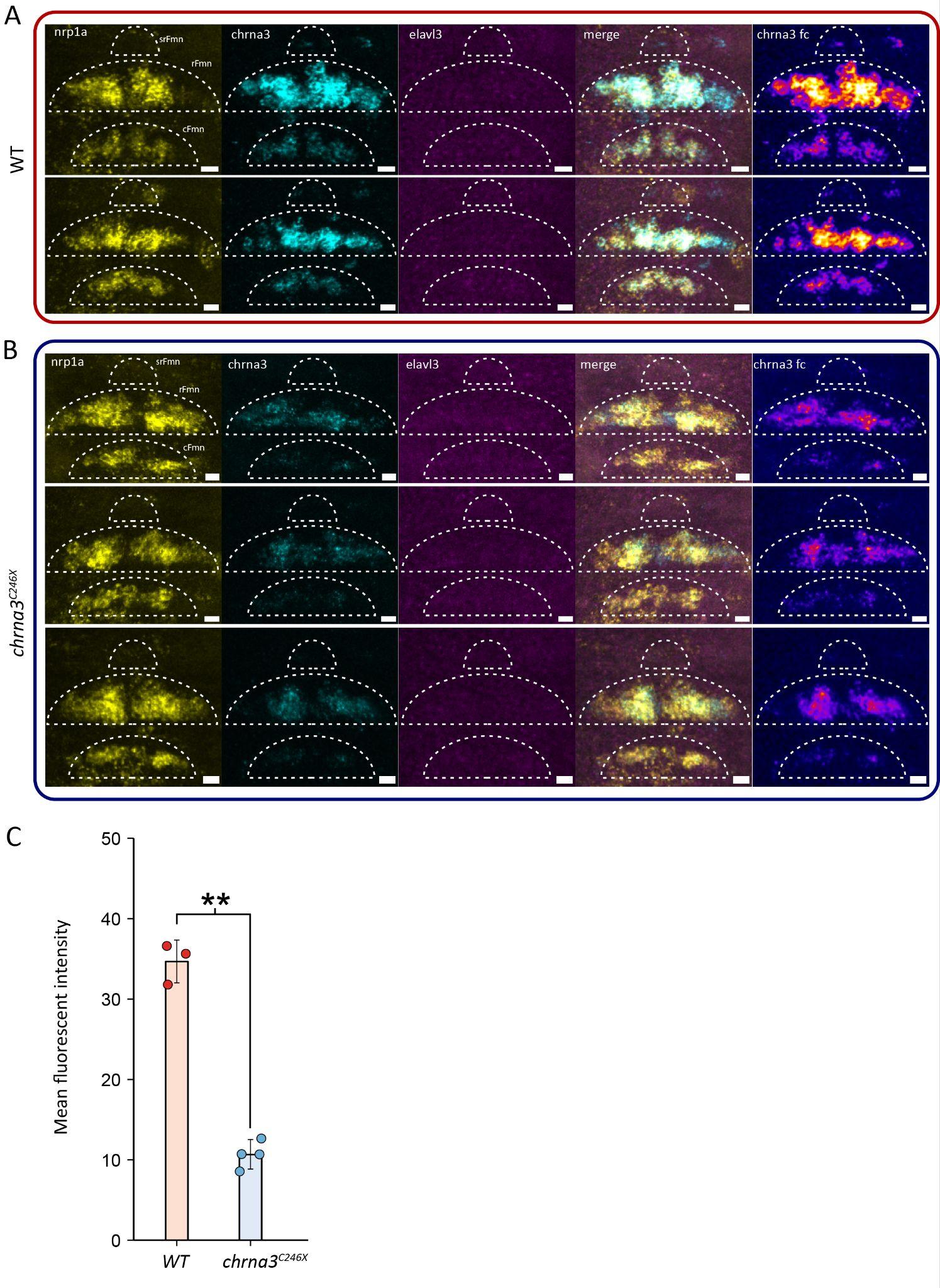

**Figure S4.** *chrna3* expression in *WT* (**A**) and *chrna3^C246X^* (**B**) 14dpf zebrafish larvae facial motor nucleus (Fmn) marked by *chata* expression, visualised by *in-situ* hybridisation chain reaction (HCR). **C.** Comparison of the mean fluorescent intensity (± standard deviation) of *chrna3* in the demarcated zone between genotypes (Mean difference = -24.0 [95CI -26.3,-20.9], p = 0.012).

Table S5. Statistical analysis output table for figure 4

| **Genotype & Metric (Figure)** | **Alcohol treatment (vs 0%)** | **Cliff's Delta** | **95% CI upper** | **95% CI lower** | **P value** |
| --- | --- | --- | --- | --- | --- |
| WT Volume dispensed (4A) | x̄0.5% | -0.554 | -0.266 | -0.759 | < 0.001 |
| WT Volume dispensed (4A) | x̄1.5% | -0.728 | -0.441 | -0.895 | < 0.001 |
| C246X Volume dispensed (4B) | x̄0.5% | -0.128 | 0.185 | -0.421 | 0.397 |
| C246X Volume dispensed (4B) | x̄1.5% | -0.583 | -0.304 | -0.791 | < 0.001 |
| WT ΔPI Pre - Stimulus 0-9m (4C) | x̄0.5% | 0.317 | 0.577 | 0.014 | 0.038 |
| WT ΔPI Pre - Stimulus 0-9m (4C) | x̄1.5% | 0.177 | 0.464 | -0.147 | 0.248 |
| C246X ΔPI Pre - Stimulus 0-9m (4D) | x̄0.5% | 0.449 | 0.682 | 0.127 | 0.002 |
| C246X ΔPI Pre - Stimulus 0-9m (4D) | x̄1.5% | 0.412 | 0.647 | 0.108 | 0.007 |
| WT ΔPI Stimulus 0-9m - 9-18m (4E) | x̄0.5% | -0.237 | 0.058 | -0.497 | 0.117 |
| WT ΔPI Stimulus 0-9m - 9-18m (4E) | x̄1.5% | -0.504 | -0.227 | -0.724 | < 0.001 |
| C246X ΔPI Stimulus 0-9m - 9-18m (4F) | x̄0.5% | -0.351 | -0.047 | -0.614 | 0.016 |
| C246X ΔPI Stimulus 0-9m - 9-18m (4F) | x̄1.5% | -0.201 | -0.0945 | -0.493 | 0.192 |

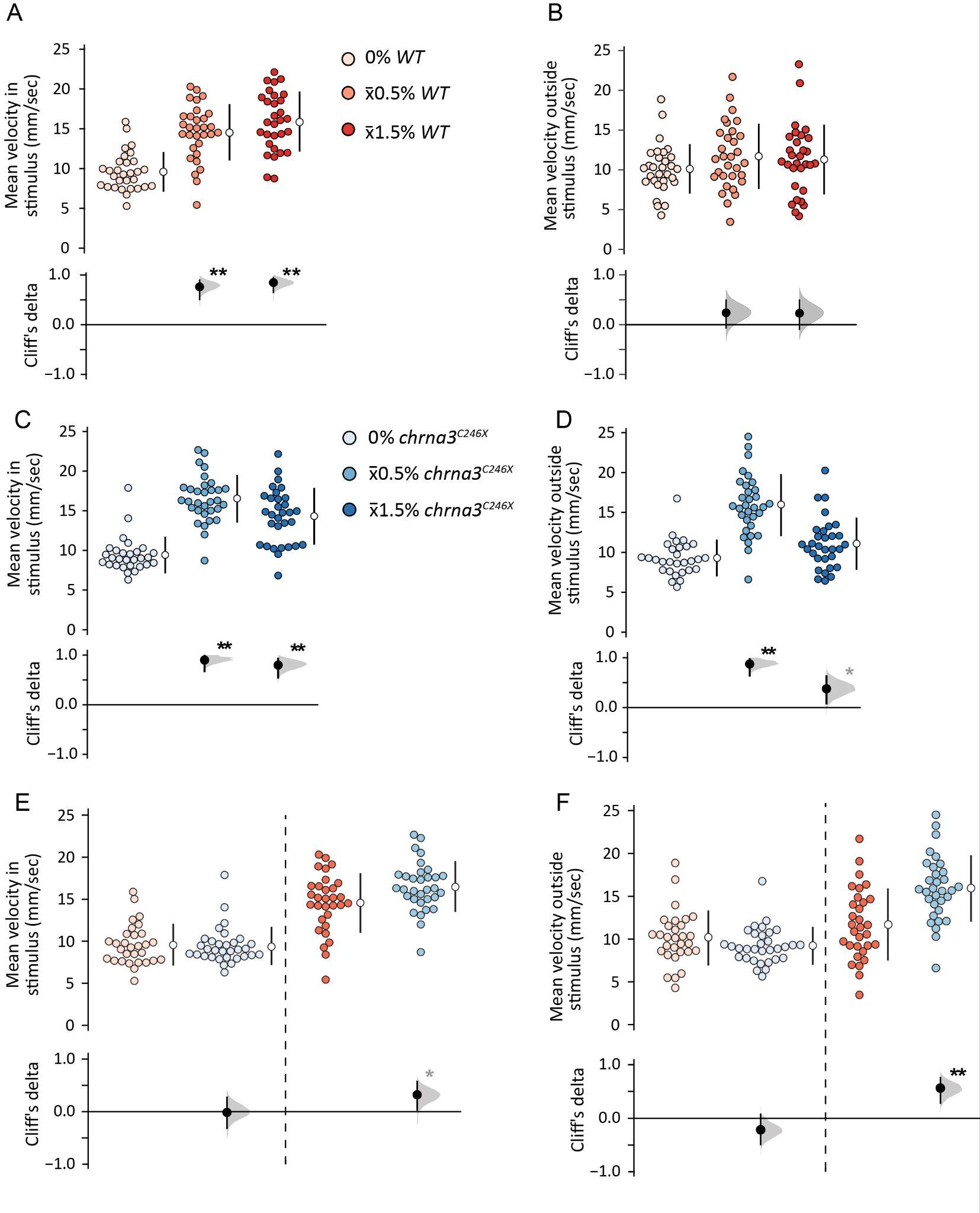

**Figure S5. *chrna3* mutant fish exhibit a stronger alcohol-induced locomotor response**

Gardener-Altman and Cumming estimation plots for total SAZA self-administration period for WT (**A, B**), *chrna3^C246X^* (**C, D**) and comparisons (**E, F**). Plots display the comparison of velocity inside (**A, C, E**) or outside (**B, D, F**) the stimulus zone at different concentrations (0/x̄0.5/x̄1.5%) for WT (n = 31/31/30) and *chrna3^C246X^* zebrafish (n = 29/31/30). Asterisks indicate the following significant differences from 0% alcohol treatments (**A-D**) or WT (**E, F**): * = p < 0.05, and effect size reported by Cliff’s delta between ± 0.2 and ± 0.4 are considered provisional, ** = p < 0.01 and effect size bigger than ± 0.4 is considered a meaningful difference. See tables S6 and S7 for exact effect size and p-values.

Table S6. Statistical analysis output table for figure S5A-D

| **Metric (Figure)** | **Alcohol treatment (vs 0%)** | **Cliff's Delta** | **95% CI upper** | **95% CI lower** | **P value** |
| --- | --- | --- | --- | --- | --- |
| WT Mean velocity in stimulus chamber (S3A) | x̄0.5% | 0.750 | 0.892 | 0.501 | < 0.001 |
| WT Mean velocity in stimulus chamber (S3A) | x̄1.5% | 0.931 | 0.98 | 0.796 | < 0.001 |
| WT Mean velocity outside stimulus chamber (S3B) | x̄0.5% | 0.239 | 0.514 | -0.067 | 0.112 |
| WT Mean velocity outside stimulus chamber (S3B) | x̄1.5% | 0.229 | 0.507 | -0.10 | 0.129 |
| Mean velocity in stimulus chamber (S3C) | x̄0.5% | 0.901 | 0.987 | 0.677 | < 0.001 |
| Mean velocity in stimulus chamber (S3C) | x̄1.5% | 0.800 | 0.927 | 0.546 | < 0.001 |
| Mean velocity outside stimulus chamber (S3D) | x̄0.5% | 0.873 | 0.972 | 0.640 | < 0.001 |
| Mean velocity outside stimulus chamber (S3D) | x̄1.5% | 0.377 | 0.626 | 0.083 | 0.011 |

Table S7. Statistical analysis output table for figure S5E-F

| **Metric (Figure)** | **Alcohol treatment (vs WT)** | **Cliff's Delta** | **95% CI upper** | **95% CI lower** | **P value** |
| --- | --- | --- | --- | --- | --- |
| Mean velocity in stimulus chamber (S3E) | 0% | -0.027 | 0.273 | -0.315 | 0.839 |
| Mean velocity in stimulus chamber (S3E) | x̄0.5% | 0.329 | 0.571 | 0.032 | 0.029 |
| Mean velocity outside stimulus chamber (S3F) | 0% | -0.217 | 0.077 | -0.488 | 0.134 |
| Mean velocity outside stimulus chamber (S3F) | x̄0.5% | 0.565 | 0.758 | 0.284 | < 0.001 |

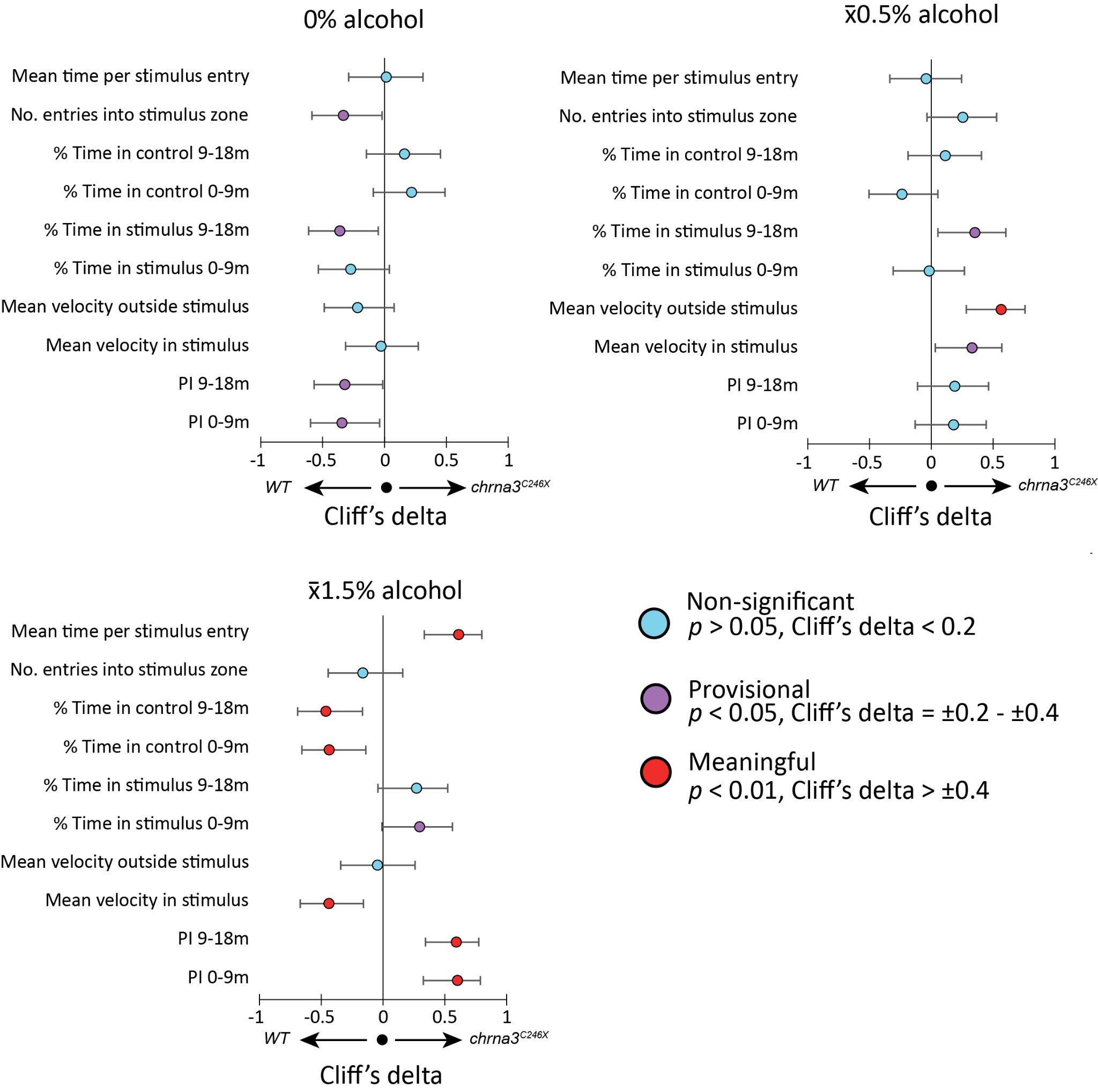

**Figure S6**. Cliff’s delta (± 95% CI) forest plots of all calculated metrics from 0/x̄0.5/x̄1.5% alcohol SAZA, comparing the response of wild type to *chrna3^C246X^* mutant zebrafish. Positive values indicate a greater response in the mutant. See table S8 for exact effect size and p-values.

Table S8. Statistical analysis output table for figure S6

| **Alcohol treatment (Figure)** | **Metric (*C246X* Vs WT)** | **Cliff's Delta** | **95% CI Lower** | **95% CI Upper** | **P value** |
| --- | --- | --- | --- | --- | --- |
| 0% (S3A) | PI 0-9 | -0.344 | -0.598 | -0.039 | 0.021 |
| 0% (S3A) | PI 9-18 | -0.320 | -0.570 | -0.014 | 0.031 |
| 0% (S3A) | Mean Vin | -0.027 | -0.315 | 0.273 | 0.8392 |
| 0% (S3A) | Mean Vout | -0.217 | -0.488 | 0.077 | 0.134 |
| 0% (S3A) | Tin Stim 0-9 | -0.274 | -0.536 | 0.039 | 0.062 |
| 0% (S3A) | Tin Stim 9-18 | -0.361 | -0.614 | -0.051 | 0.013 |
| 0% (S3A) | Tin control 0-9 | 0.218 | -0.092 | 0.490 | 0.147 |
| 0% (S3A) | Tin control 9-18 | 0.161 | -0.147 | 0.453 | 0.270 |
| 0% (S3A) | Entries into stim | -0.332 | -0.586 | -0.020 | 0.026 |
| 0% (S3A) | Mean time per stim entry | 0.014 | -0.290 | 0.310 | 0.922 |
| x̄0.5% (S3B) | PI 0-9 | 0.180 | -0.130 | 0.444 | 0.231 |
| x̄0.5% (S3B) | PI 9-18 | 0.190 | -0.111 | 0.464 | 0.199 |
| x̄0.5% (S3B) | Mean Vin | 0.329 | 0.032 | 0.571 | 0.029 |
| x̄0.5% (S3B) | Mean Vout | 0.565 | 0.284 | 0.758 | 0.000 |
| x̄0.5% (S3B) | Tin Stim 0-9 | -0.016 | -0.308 | 0.268 | 0.910 |
| x̄0.5% (S3B) | Tin Stim 9-18 | 0.353 | 0.054 | 0.603 | 0.015 |
| x̄0.5% (S3B) | Tin control 0-9 | -0.238 | -0.504 | 0.054 | 0.102 |
| x̄0.5% (S3B) | Tin control 9-18 | 0.113 | -0.188 | 0.407 | 0.454 |
| x̄0.5% (S3B) | Entries into stim | 0.255 | -0.034 | 0.528 | 0.085 |
| x̄0.5% (S3B) | Mean time per stim entry | -0.042 | -0.335 | 0.246 | 0.772 |
| x̄1.5% (S3C) | PI 0-9 | 0.602 | 0.325 | 0.787 | 0.000 |
| x̄1.5% (S3C) | PI 9-18 | 0.594 | 0.342 | 0.774 | 0.000 |
| x̄1.5% (S3C) | Mean Vin | -0.4370 | -0.6710 | -0.1590 | 0.002 |
| x̄1.5% (S3C) | Mean Vout | -0.0450 | -0.3440 | 0.2580 | 0.761 |
| x̄1.5% (S3C) | Tin Stim 0-9 | 0.2970 | -0.0090 | 0.5610 | 0.0466 |
| x̄1.5% (S3C) | Tin Stim 9-18 | 0.2710 | -0.0430 | 0.5230 | 0.0658 |
| x̄1.5% (S3C) | Tin control 0-9 | -0.434 | -0.658 | -0.140 | 0.004 |
| x̄1.5% (S3C) | Tin control 9-18 | -0.462 | -0.691 | -0.168 | 0.001 |
| x̄1.5% (S3C) | Entries into stim | -0.163 | -0.444 | 0.158 | 0.280 |
| x̄1.5% (S3C) | Mean time per stim entry | 0.613 | 0.333 | 0.798 | 0.000 |

Table S9. Statistical analysis output table for figure 5A-G

| **Genotype & Metric (Figure)** | **Alcohol treatment** | **Time period vs 0% (m)** | **Cliff's Delta** | **95% CI upper** | **95% CI lower** | **P value** |
| --- | --- | --- | --- | --- | --- | --- |
| WT Total area between fish (5B) | 0.5% | 0-30 | 0.409 | 0.453 | 0.364 | < 0.001 |
| WT Total area between fish (5B) | 1% | 0-30 | 0.518 | 0.559 | 0.475 | < 0.001 |
| C246X Total area between fish (5C) | 0.5% | 0-30 | 0.365 | 0.412 | 0.320 | < 0.001 |
| C246X Total area between fish (5C) | 1% | 0-30 | 0.611 | 0.649 | 0.573 | < 0.001 |
| WT 18X3m Total area (5D) | 0.5% | 0-3 | 0.147 | 0.18 | 0.111 | < 0.001 |
| WT 18X3m Total area (5D) | 1% | 0-3 | 0.225 | 0.258 | 0.19 | < 0.001 |
| WT 18X3m Total area (5D) | 0.5% | 3-6 | 0.287 | 0.32 | 0.252 | < 0.001 |
| WT 18X3m Total area (5D) | 1% | 3-6 | 0.338 | 0.37 | 0.305 | < 0.001 |
| WT 18X3m Total area (5D) | 0.5% | 6-9 | 0.335 | 0.368 | 0.303 | < 0.001 |
| WT 18X3m Total area (5D) | 1% | 6-9 | 0.417 | 0.447 | 0.386 | < 0.001 |
| WT 18X3m Total area (5D) | 0.5% | 9-12 | 0.455 | 0.486 | 0.425 | < 0.001 |
| WT 18X3m Total area (5D) | 1% | 9-12 | 0.578 | 0.605 | 0.551 | < 0.001 |
| WT 18X3m Total area (5D) | 0.5% | 12-15 | 0.552 | 0.579 | 0.522 | < 0.001 |
| WT 18X3m Total area (5D) | 1% | 12-15 | 0. 728 | 0.749 | 0.707 | < 0.001 |
| WT 18X3m Total area (5D) | 0.5% | 15-18 | 0.358 | 0.389 | 0.325 | < 0.001 |
| WT 18X3m Total area (5D) | 1% | 15-18 | 0. 284 | 0.316 | 0.249 | < 0.001 |
| C246X 18x3m Total area (5E) | 0.5% | 0-3 | 0.327 | 0.36 | 0.293 | < 0.001 |
| C246X 18x3m Total area (5E) | 1% | 0-3 | 0.188 | 0.223 | 0.154 | < 0.001 |
| C246X 18x3m Total area (5E) | 0.5% | 3-6 | 0.321 | 0.354 | 0.287 | < 0.001 |
| C246X 18x3m Total area (5E) | 1% | 3-6 | 0.474 | 0.503 | 0.442 | < 0.001 |
| C246X 18x3m Total area (5E) | 0.5% | 6-9 | 0.386 | 0.418 | 0.353 | < 0.001 |
| C246X 18x3m Total area (5E) | 1% | 6-9 | 0.598 | 0.625 | 0.571 | < 0.001 |
| C246X 18x3m Total area (5E) | 0.5% | 9-12 | 0.277 | 0.311 | 0.243 | < 0.001 |
| C246X 18x3m Total area (5E) | 1% | 9-12 | 0.536 | 0.565 | 0.507 | < 0.001 |
| C246X 18x3m Total area (5E) | 0.5% | 12-15 | 0.329 | 0.362 | 0.295 | < 0.001 |
| C246X 18x3m Total area (5E) | 1% | 12-15 | 0.664 | 0.688 | 0.639 | < 0.001 |
| C246X 18x3m Total area (5E) | 0.5% | 15-18 | 0.326 | 0.358 | 0.291 | < 0.001 |
| C246X 18x3m Total area (5E) | 1% | 15-18 | 0.782 | 0.8 | 0.76 | < 0.001 |
| WT 18x3m Kinesis (5F) | 0.5% | 0-3 | -0.047 | -0.013 | -0.084 | 0.01 |
| WT 18x3m Kinesis (5F) | 1% | 0-3 | -0.288 | -0.255 | -0.32 | < 0.001 |
| WT 18x3m Kinesis (5F) | 0.5% | 3-6 | -0.028 | 0.007 | -0.063 | 0.116 |
| WT 18x3m Kinesis (5F) | 1% | 3-6 | -0.322 | -0.289 | -0.353 | < 0.001 |
| WT 18x3m Kinesis (5F) | 0.5% | 6-9 | -0.024 | 0.011 | -0.061 | 0.189 |
| WT 18x3m Kinesis (5F) | 1% | 6-9 | -0.419 | -0.387 | -0.449 | < 0.001 |
| WT 18x3m Kinesis (5F) | 0.5% | 9-12 | -0.014 | 0.022 | -0.049 | 0.423 |
| WT 18x3m Kinesis (5F) | 1% | 9-12 | -0.422 | -0.391 | -0.451 | <0.001 |
| WT 18x3m Kinesis (5F) | 0.5% | 12-15 | 0.069 | 0.104 | 0.035 | < 0.001 |
| WT 18x3m Kinesis (5F) | 1% | 12-15 | -0.402 | -0.37 | -0.434 | < 0.001 |
| WT 18x3m Kinesis (5F) | 0.5% | 15-18 | 0.083 | 0.117 | 0.049 | < 0.001 |
| WT 18x3m Kinesis (5F) | 1% | 15-18 | -0.408 | -0.375 | -0.44 | < 0.001 |
| C246X 18x3m Kinesis (5G) | 0.5% | 0-3 | 0.113 | 0.149 | 0.077 | < 0.001 |
| C246X 18x3m Kinesis (5G) | 1% | 0-3 | -0.204 | -0.171 | -0.24 | < 0.001 |
| C246X 18x3m Kinesis (5G) | 0.5% | 3-6 | 0.108 | 0.146 | 0.074 | < 0.001 |
| C246X 18x3m Kinesis (5G) | 1% | 3-6 | -0.296 | -0.26 | -0.33 | < 0.001 |
| C246X 18x3m Kinesis (5G) | 0.5% | 6-9 | 0.03 | 0.066 | -0.007 | 0.113 |
| C246X 18x3m Kinesis (5G) | 1% | 6-9 | -0.243 | 0.208 | 0.277 | < 0.001 |
| C246X 18x3m Kinesis (5G) | 0.5% | 9-12 | -0.047 | -0.009 | -0.083 | 0.011 |
| C246X 18x3m Kinesis (5G) | 1% | 9-12 | -0.195 | -0.16 | -0.23 | < 0.001 |
| C246X 18x3m Kinesis (5G) | 0.5% | 12-15 | -0.035 | 0.001 | -0.071 | 0.061 |
| C246X 18x3m Kinesis (5G) | 1% | 12-15 | -0.203 | -0.167 | -0.238 | < 0.001 |
| C246X 18x3m Kinesis (5G) | 0.5% | 15-18 | -0.074 | -0.039 | -0.11 | < 0.001 |
| C246X 18x3m Kinesis (5G) | 1% | 15-18 | -0.067 | -0.031 | -0.102 | < 0.001 |

Table S10. Statistical analysis output table for figure 5H-I

| **Assay** **(Figure)** | **Time point (*C246X* vs WT)** | **Cliff's Delta** | **95% CI upper** | **95% CI lower** | **P value** |
| --- | --- | --- | --- | --- | --- |
| 0.5% alcohol 18X3m total area (5H) | 0-3m | 0.027 | 0.062 | -0.009 | 0.131 |
| 0.5% alcohol 18X3m total area (5H) | 3-6m | -0.156 | -0.122 | -0.192 | < 0.001 |
| 0.5% alcohol 18X3m total area (5H) | 6-9m | -0.223 | -0.19 | -0.256 | < 0.001 |
| 0.5% alcohol 18X3m total area (5H) | 9-12m | -0.283 | -0.25 | -0.317 | < 0.001 |
| 0.5% alcohol 18X3m total area (5H) | 12-15m | -0.44 | -0.409 | -0.47 | < 0.001 |
| 0.5% alcohol 18X3m total area (5H) | 15-18m | -0.356 | -0.323 | -0.387 | < 0.001 |
| 0.5% alcohol 18X3m Kinesis (5I) | 0-3m | 0.253 | 0.288 | 0.219 | < 0.001 |
| 0.5% alcohol 18X3m Kinesis (5I) | 3-6m | 0.177 | 0.214 | 0.142 | < 0.001 |
| 0.5% alcohol 18X3m Kinesis (5I) | 6-9m | 0.105 | 0.142 | 0.071 | < 0.001 |
| 0.5% alcohol 18X3m Kinesis (5I) | 9-12m | -0.017 | 0.016 | -0.054 | 0.338 |
| 0.5% alcohol 18X3m Kinesis (5I) | 12-15m | -0.026 | 0.009 | -0.062 | 0.143 |
| 0.5% alcohol 18X3m Kinesis (5I) | 15-18m | -0.069 | -0.034 | -0.105 | < 0.001 |

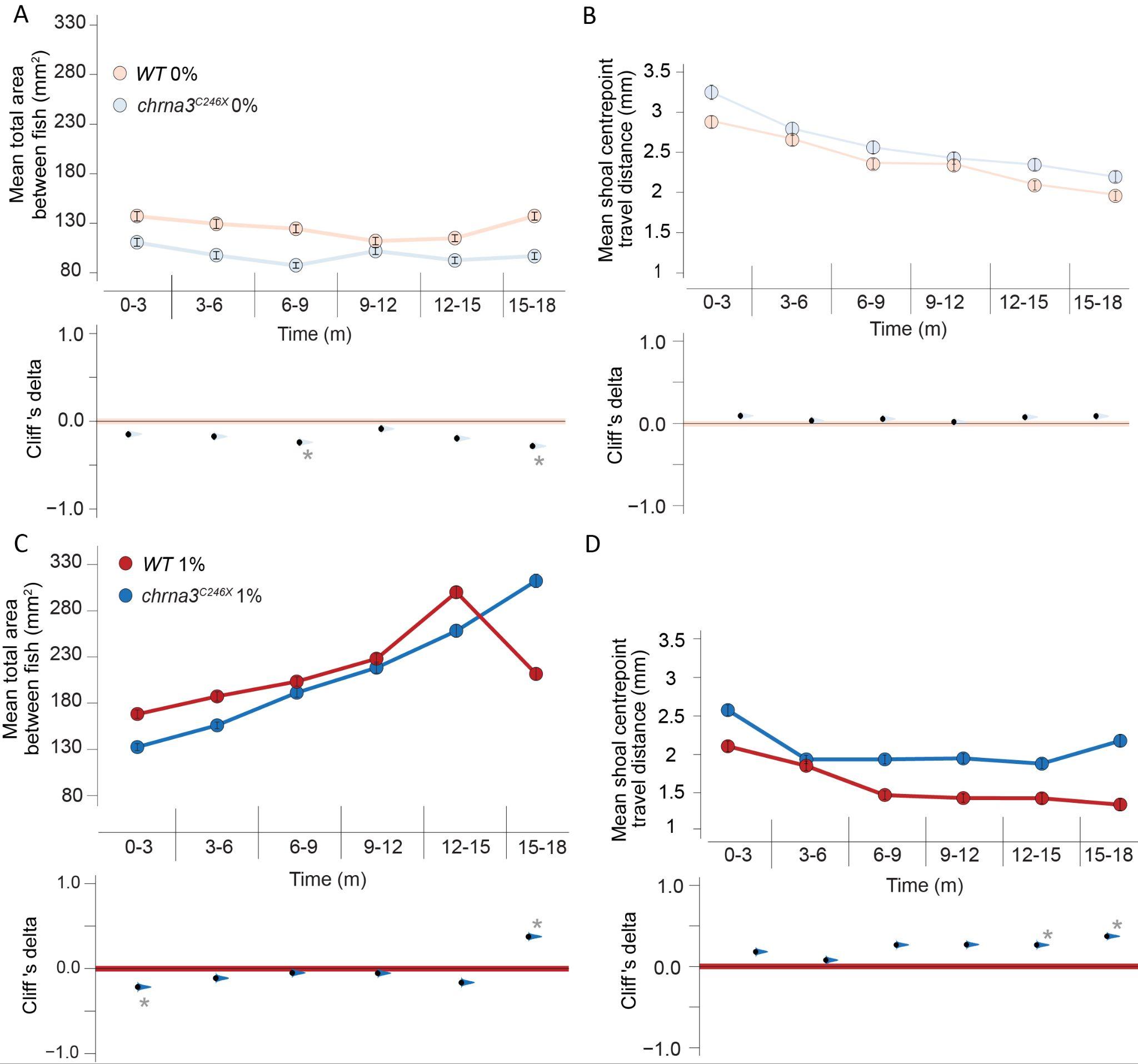

**Figure S7**. ***WT* and *chrna3^C246X^*** **mutants share shoaling cohesion and kinesis phenotypes during 0% and 1% alcohol immersion.** Mean (± 95% CI) shoal cohesion (**A, C**) and kinesis (**B, D**) with Cumming estimation plots comparing 0% (**A, B**), or 1% alcohol exposure (**C, D**) between genotypes. Metrics compared per frame at 2fps (n = 6 assays per genotype, per treatment). Asterisks indicate the following significant differences from WT within that time period: * = p < 0.05, and effect size reported by Cliff’s delta between ± 0.2 and ± 0.4 are considered provisional, ** = p < 0.01 and effect size bigger than ± 0.4 is considered a meaningful difference. See table S11 for exact effect size and p-values.

Table S11. Statistical analysis output table for figure S7

| **Assay** **(Figure)** | **Time point (*C246X* vs WT)** | **Cliff's Delta** | **95% CI upper** | **95% CI lower** | **P value** |
| --- | --- | --- | --- | --- | --- |
| 0% alcohol 18X3m total area (S5A) | 0-3m | -0.147 | -0.112 | -0.183 | < 0.001 |
| 0% alcohol 18X3m total area (S5A) | 3-6m | -0.174 | -0.138 | -0.209 | < 0.001 |
| 0% alcohol 18X3m total area (S5A) | 6-9m | -0.24 | -0.206 | -0.276 | < 0.001 |
| 0% alcohol 18X3m total area (S5A) | 9-12m | -0.086 | -0.048 | -0.12 | < 0.001 |
| 0% alcohol 18X3m total area (S5A) | 12-15m | -0.195 | -0.16 | -0.229 | < 0.001 |
| 0% alcohol 18X3m total area (S5A) | 15-18m | -0.283 | -0.249 | -0.317 | < 0.001 |
| 0% alcohol 18X3m Kinesis (S5B) | 0-3m | 0.092 | 0.128 | 0.055 | < 0.001 |
| 0% alcohol 18X3m Kinesis (S5B) | 3-6m | 0.033 | 0.071 | -0.003 | 0.079 |
| 0% alcohol 18X3m Kinesis (S5B) | 6-9m | 0.055 | 0.091 | 0.017 | 0.003 |
| 0% alcohol 18X3m Kinesis (S5B) | 9-12m | 0.016 | 0.052 | -0.021 | 0.383 |
| 0% alcohol 18X3m Kinesis (S5B) | 12-15m | 0.076 | 0.111 | 0.04 | < 0.001 |
| 0% alcohol 18X3m Kinesis (S5B) | 15-18m | 0.088 | 0.124 | 0.053 | < 0.001 |
| 1% alcohol 18X3m total area (S5C) | 0-3m | -0.216 | -0.18 | -0.247 | < 0.001 |
| 1% alcohol 18X3m total area (S5C) | 3-6m | -0.112 | -0.078 | -0.147 | < 0.001 |
| 1% alcohol 18X3m total area (S5C) | 6-9m | -0.049 | -0.015 | -0.082 | < 0.001 |
| 1% alcohol 18X3m total area (S5C) | 9-12m | -0.053 | -0.018 | -0.087 | 0.003 |
| 1% alcohol 18X3m total area (S5C) | 12-15m | -0.163 | -0.129 | -0.199 | < 0.001 |
| 1% alcohol 18X3m total area (S5C) | 15-18m | 0.377 | 0.41 | 0.345 | < 0.001 |
| 1% alcohol 18X3m Kinesis (S5D) | 0-3m | 0.181 | 0.213 | 0.145 | < 0.001 |
| 1% alcohol 18X3m Kinesis (S5D) | 3-6m | 0.078 | 0.113 | 0.042 | < 0.001 |
| 1% alcohol 18X3m Kinesis (S5D) | 6-9m | 0.266 | 0.299 | 0.232 | < 0.001 |
| 1% alcohol 18X3m Kinesis (S5D) | 9-12m | 0.27 | 0.302 | 0.235 | < 0.001 |
| 1% alcohol 18X3m Kinesis (S5D) | 12-15m | 0.265 | 0.298 | 0.233 | < 0.001 |
| 1% alcohol 18X3m Kinesis (S5D) | 15-18m | 0.372 | 0.405 | 0.34 | < 0.001 |

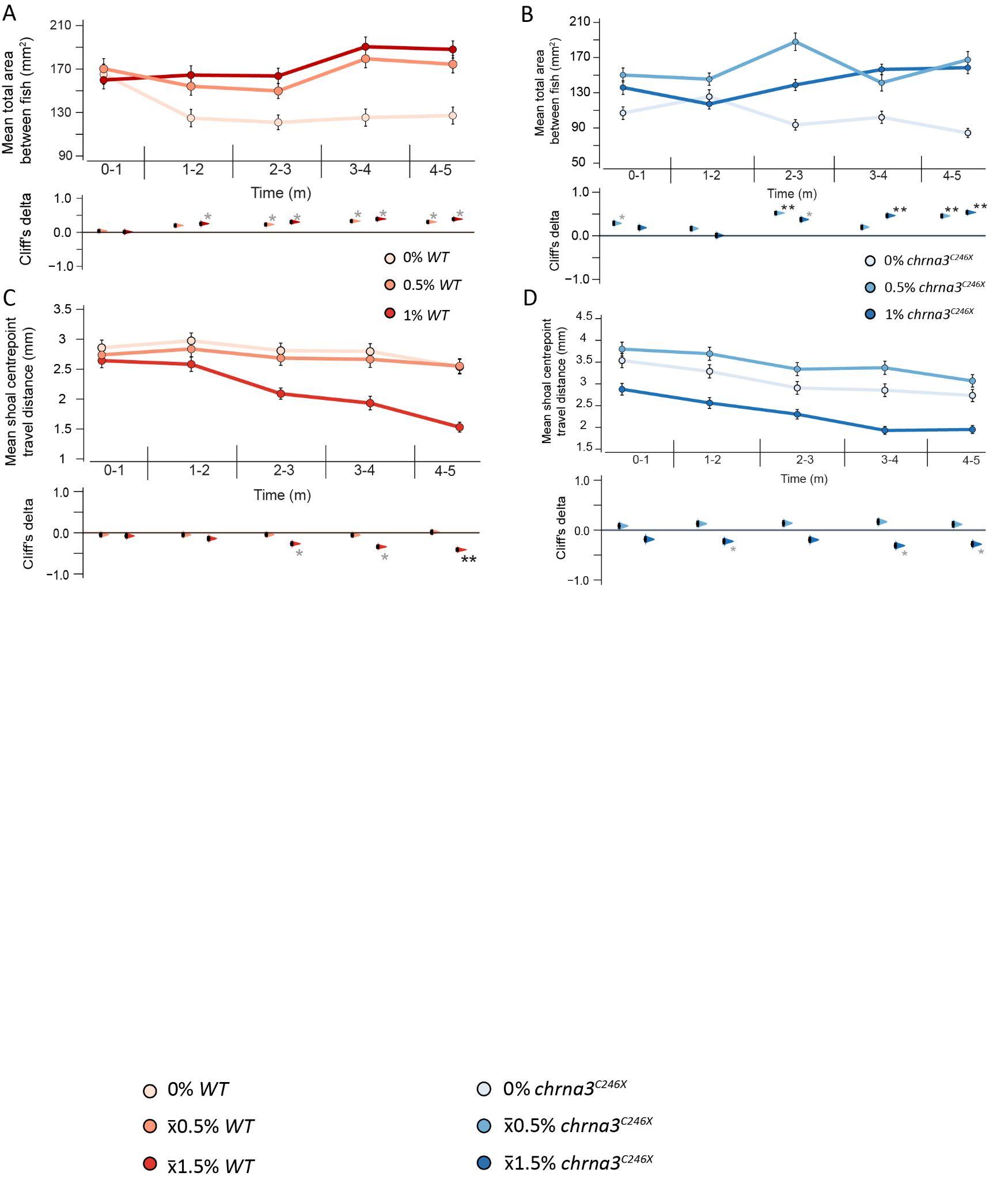

**Figure S8. *WT* and *chrna3^C246X^* zebrafish exhibit shoal cohesion and kinesis change within five minutes of alcohol immersion.** Mean (± 95% CI) shoal cohesion (**A, B**) and kinesis (**C, D**) with Cumming estimation plots in WT (**A, C**), or *chrna3^C246X^* (**B, D**) under the first five minutes of 0/0.5/1% alcohol exposure. Metrics compared per frame at 2fps (n = 6 assays per genotype, per treatment). Asterisks indicate the following significant differences from 0% within that exposure concentration: * = p < 0.05, and effect size reported by Cliff’s delta between ± 0.2 and ± 0.4 are considered provisional, ** = p < 0.01 and effect size bigger than ± 0.4 is considered a meaningful difference. See table S12 for exact effect size and p-values.

Table S12. Statistical analysis output table for figure S8

| **Metric (Figure)** | **Alcohol treatment** | **Time period vs 0% (m)** | **Cliff's Delta** | **95% CI upper** | **95% CI lower** | **P value** |
| --- | --- | --- | --- | --- | --- | --- |
| WT 5X1m Total area (S6A) | 0.5% | 0-1 | 0.0314 | 0.0891 | -0.0304 | 0.383 |
| WT 5X1m Total area (S6A) | 1% | 0-1 | 0.0012 | 0.0592 | -0.059 | 0.363 |
| WT 5X1m Total area (S6A) | 0.5% | 1-2 | 0.194 | 0.253 | 0.129 | < 0.001 |
| WT 5X1m Total area (S6A) | 1% | 1-2 | 0.247 | 0.306 | 0.185 | < 0.001 |
| WT 5X1m Total area (S6A) | 0.5% | 2-3 | 0.224 | 0.282 | 0.165 | < 0.001 |
| WT 5X1m Total area (S6A) | 1% | 2-3 | 0.298 | 0.356 | 0.241 | < 0.001 |
| WT 5X1m Total area (S6A) | 0.5% | 3-4 | 0.324 | 0.382 | 0.266 | < 0.001 |
| WT 5X1m Total area (S6A) | 1% | 3-4 | 0.387 | 0.443 | 0.331 | < 0.001 |
| WT 5X1m Total area (S6A) | 0.5% | 4-5 | 0.3 | 0.358 | 0.242 | < 0.001 |
| WT 5X1m Total area (S6A) | 1% | 4-5 | 0.383 | 0.437 | 0.328 | < 0.001 |
| C246X 5X1m Total area (S6B) | 0.5% | 0-1 | 0.327 | 0.36 | 0.293 | < 0.001 |
| C246X 5X1m Total area (S6B) | 1% | 0-1 | 0.188 | 0.223 | 0.154 | < 0.001 |
| C246X 5X1m Total area (S6B) | 0.5% | 1-2 | 0.321 | 0.354 | 0.287 | < 0.001 |
| C246X 5X1m Total area (S6B) | 1% | 1-2 | 0.474 | 0.503 | 0.442 | < 0.001 |
| C246X 5X1m Total area (S6B) | 0.5% | 2-3 | 0.386 | 0.418 | 0.353 | < 0.001 |
| C246X 5X1m Total area (S6B) | 1% | 2-3 | 0.598 | 0.625 | 0.571 | < 0.001 |
| C246X 5X1m Total area (S6B) | 0.5% | 3-4 | 0.277 | 0.311 | 0.243 | < 0.001 |
| C246X 5X1m Total area (S6B) | 1% | 3-4 | 0.536 | 0.565 | 0.507 | < 0.001 |
| C246X 5X1m Total area (S6B) | 0.5% | 4-5 | 0.329 | 0.362 | 0.295 | < 0.001 |
| C246X 5X1m Total area (S6B) | 1% | 4-5 | 0.664 | 0.688 | 0.639 | < 0.001 |
| WT 5X1m Kinesis (S6C) | 0.5% | 0-1 | -0.0536 | 0.0106 | -0.114 | 0.111 |
| WT 5X1m Kinesis (S6C) | 1% | 0-1 | -0.0747 | -0.0142 | -0.136 | 0.015 |
| WT 5X1m Kinesis (S6C) | 0.5% | 1-2 | -0.0729 | -0.0104 | -0.133 | 0.017 |
| WT 5X1m Kinesis (S6C) | 1% | 1-2 | -0.136 | -0.0747 | -0.197 | < 0.001 |
| WT 5X1m Kinesis (S6C) | 0.5% | 2-3 | -0.055 | 0.00798 | -0.114 | 0.082 |
| WT 5X1m Kinesis (S6C) | 1% | 2-3 | -0.262 | -0.202 | -0.318 | < 0.001 |
| WT 5X1m Kinesis (S6C) | 0.5% | 3-4 | -0.0592 | 0.00201 | -0.12 | 0.099 |
| WT 5X1m Kinesis (S6C) | 1% | 3-4 | -0.327 | -0.271 | -0.383 | < 0.001 |
| WT 5X1m Kinesis (S6C) | 0.5% | 4-5 | 0.0138 | 0.0745 | -0.0476 | 0.957 |
| WT 5X1m Kinesis (S6C) | 1% | 4-5 | -0.401 | -0.345 | -0.453 | < 0.001 |
| C246X 5X1m Kinesis (S6D) | 0.5% | 0-1 | 0.113 | 0.149 | 0.077 | < 0.001 |
| C246X 5X1m Kinesis (S6D) | 1% | 0-1 | -0.204 | -0.171 | -0.24 | < 0.001 |
| C246X 5X1m Kinesis (S6D) | 0.5% | 1-2 | 0.108 | 0.146 | 0.074 | < 0.001 |
| C246X 5X1m Kinesis (S6D) | 1% | 1-2 | -0.296 | -0.26 | -0.33 | < 0.001 |
| C246X 5X1m Kinesis (S6D) | 0.5% | 2-3 | 0.03 | 0.066 | -0.007 | 0.113 |
| C246X 5X1m Kinesis (S6D) | 1% | 2-3 | -0.243 | 0.208 | 0.277 | < 0.001 |
| C246X 5X1m Kinesis (S6D) | 0.5% | 3-4 | -0.047 | -0.009 | -0.083 | 0.011 |
| C246X 5X1m Kinesis (S6D) | 1% | 3-4 | -0.195 | -0.16 | -0.23 | < 0.001 |
| C246X 5X1m Kinesis (S6D) | 0.5% | 4-5 | -0.035 | 0.001 | -0.071 | 0.061 |
| C246X 5X1m Kinesis (S6D) | 1% | 4-5 | -0.203 | -0.167 | -0.238 | < 0.001 |

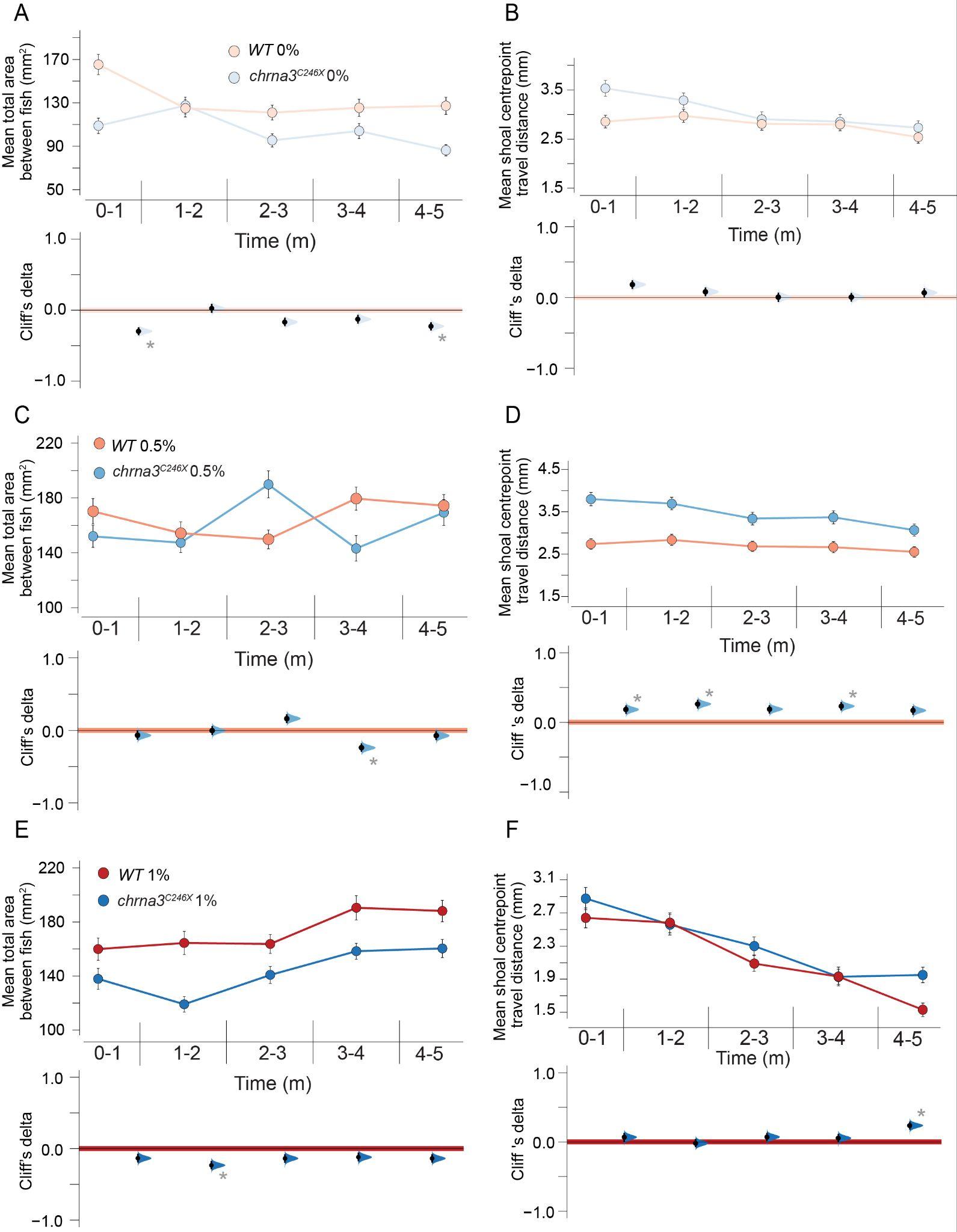

**Figure S9. *WT* and *chrna3^C246X^* zebrafish exhibit similar changes to shoal cohesion and kinesis in the first five minutes of alcohol immersion.** Mean (± 95% CI) shoal cohesion (**A, C, E**) and kinesis (**B, D, F**) with Cumming estimation plots comparing between genotypes in first five minutes of 0% (**A, B**), 0.5% (**C, D**) or 1% alcohol exposure (**E, F**). Metrics compared per frame at 2fps (n = 6 assays per genotype, per treatment). Asterisks indicate the following significant differences from WT within that time period: * = p < 0.05, and effect size reported by Cliff’s delta between ± 0.2 and ± 0.4 are considered provisional, ** = p < 0.01 and effect size bigger than ± 0.4 is considered a meaningful difference. See table S13 for exact effect size and p-values.

Table S13. Statistical analysis output table for figure S9

| **Assay** **(Figure)** | **Time point (*C246X* vs WT)** | **Cliff's Delta** | **95% CI upper** | **95% CI lower** | **P value** |
| --- | --- | --- | --- | --- | --- |
| 0% alcohol 5X1m total area (S7A) | 0-1m | -0.297 | -0.24 | -0.353 | < 0.001 |
| 0% alcohol 5X1m total area (S7A) | 1-2m | 0.026 | 0.088 | -0.038 | 0.407 |
| 0% alcohol 5X1m total area (S7A) | 2-3m | -0.168 | -0.105 | -0.228 | < 0.001 |
| 0% alcohol 5X1m total area (S7A) | 3-4m | -0.127 | -0.065 | -0.188 | < 0.001 |
| 0% alcohol 5X1m total area (S7A) | 4-5m | -0.227 | -0.169 | -0.287 | < 0.001 |
| 0% alcohol 5X1m Kinesis (S7B) | 0-1m | 0.183 | 0.243 | 0.121 | < 0.001 |
| 0% alcohol 5X1m Kinesis (S7B) | 1-2m | 0.081 | 0.142 | 0.018 | 0.01 |
| 0% alcohol 5X1m Kinesis (S7B) | 2-3m | 0.005 | 0.068 | -0.059 | 0.866 |
| 0% alcohol 5X1m Kinesis (S7B) | 3-4m | 0.006 | 0.065 | -0.058 | 0.859 |
| 0% alcohol 5X1m Kinesis (S7B) | 4-5m | 0.067 | 0.131 | 0.004 | 0.038 |
| 0.5% alcohol 5X1m total area (S7C) | 0-1m | -0.067 | -0.005 | -0.127 | 0.029 |
| 0.5% alcohol 5X1m total area (S7C) | 1-2m | -0.003 | 0.061 | -0.063 | 0.934 |
| 0.5% alcohol 5X1m total area (S7C) | 2-3m | 0.163 | 0.223 | 0.101 | < 0.001 |
| 0.5% alcohol 5X1m total area (S7C) | 3-4m | -0.239 | -0.177 | -0.297 | < 0.001 |
| 0.5% alcohol 5X1m total area (S7C) | 4-5m | -0.073 | -0.011 | -0.134 | 0.02 |
| 0.5% alcohol 5X1m Kinesis (S7D) | 0-1m | 0.309 | 0.366 | 0.251 | < 0.001 |
| 0.5% alcohol 5X1m Kinesis (S7D) | 1-2m | 0.262 | 0.323 | 0.2 | < 0.001 |
| 0.5% alcohol 5X1m Kinesis (S7D) | 2-3m | 0.188 | 0.247 | 0.127 | < 0.001 |
| 0.5% alcohol 5X1m Kinesis (S7D) | 3-4m | 0.231 | 0.289 | 0.169 | < 0.001 |
| 0.5% alcohol 5X1m Kinesis (S7D) | 4-5m | 0.17 | 0.231 | 0.109 | < 0.001 |
| 1% alcohol 5X1m total area (S7E) | 0-1m | -0.135 | -0.073 | -0.193 | < 0.001 |
| 1% alcohol 5X1m total area (S7E) | 1-2m | -0.233 | -0.174 | -0.289 | < 0.001 |
| 1% alcohol 5X1m total area (S7E) | 2-3m | -0.137 | -0.078 | -0.198 | < 0.001 |
| 1% alcohol 5X1m total area (S7E) | 3-4m | -0.12 | -0.058 | -0.178 | < 0.001 |
| 1% alcohol 5X1m total area (S7E) | 4-5m | -0.137 | -0.078 | -0.196 | < 0.001 |
| 1% alcohol 5X1m Kinesis (S7F) | 0-1m | 0.067 | 0.128 | 0.001 | 0.032 |
| 1% alcohol 5X1m Kinesis (S7F) | 1-2m | -0.02 | 0.043 | -0.077 | 0.526 |
| 1% alcohol 5X1m Kinesis (S7F) | 2-3m | 0.07 | 0.128 | 0.009 | 0.023 |
| 1% alcohol 5X1m Kinesis (S7F) | 3-4m | 0.054 | 0.113 | -0.006 | 0.081 |
| 1% alcohol 5X1m Kinesis (S7F) | 4-5m | 0.235 | 0.292 | 0.177 | < 0.001 |

Table S14. Genes included in the custom sets ‘Nicotine dependence ’ and ‘Alcohol dependence’. These genes have been associated with nicotine and alcohol dependence, described by Hu *et al.* (Hu et al. 2018)

| **Gene set** | **Gene list** |
| --- | --- |
| Nicotine dependence | grin1a  gabra4  grin2aa  grin2bb  GRIN2B  GABRA2A  chrnb2  chrnb5b  chrna6  chrna7  FO907089.1 |
| Alcohol dependence | camk4  ppp1r1b  ddc  gnas  grin3a  npy  bdnf  grin2b  shc3  th  slc6a3  slc18a2 |
| Mammalian anxiety phenotypes | kctd10  mmab  ube3b  mapt  kansl1  tcea2  si:dkey-215k6.1  pde4bb glrbb  rbfox  tmem106ba  esr1  ntrk2b  slc6a4a  mao  comta  htr1aa  htr1b  htr2aa  htr1b  htr2aa  htr2cl1  pde4ba  glrba  tmem106bb  ntrk2a  slc6a4b  comtb  htr1ab  htr2ab  slc6a3 |

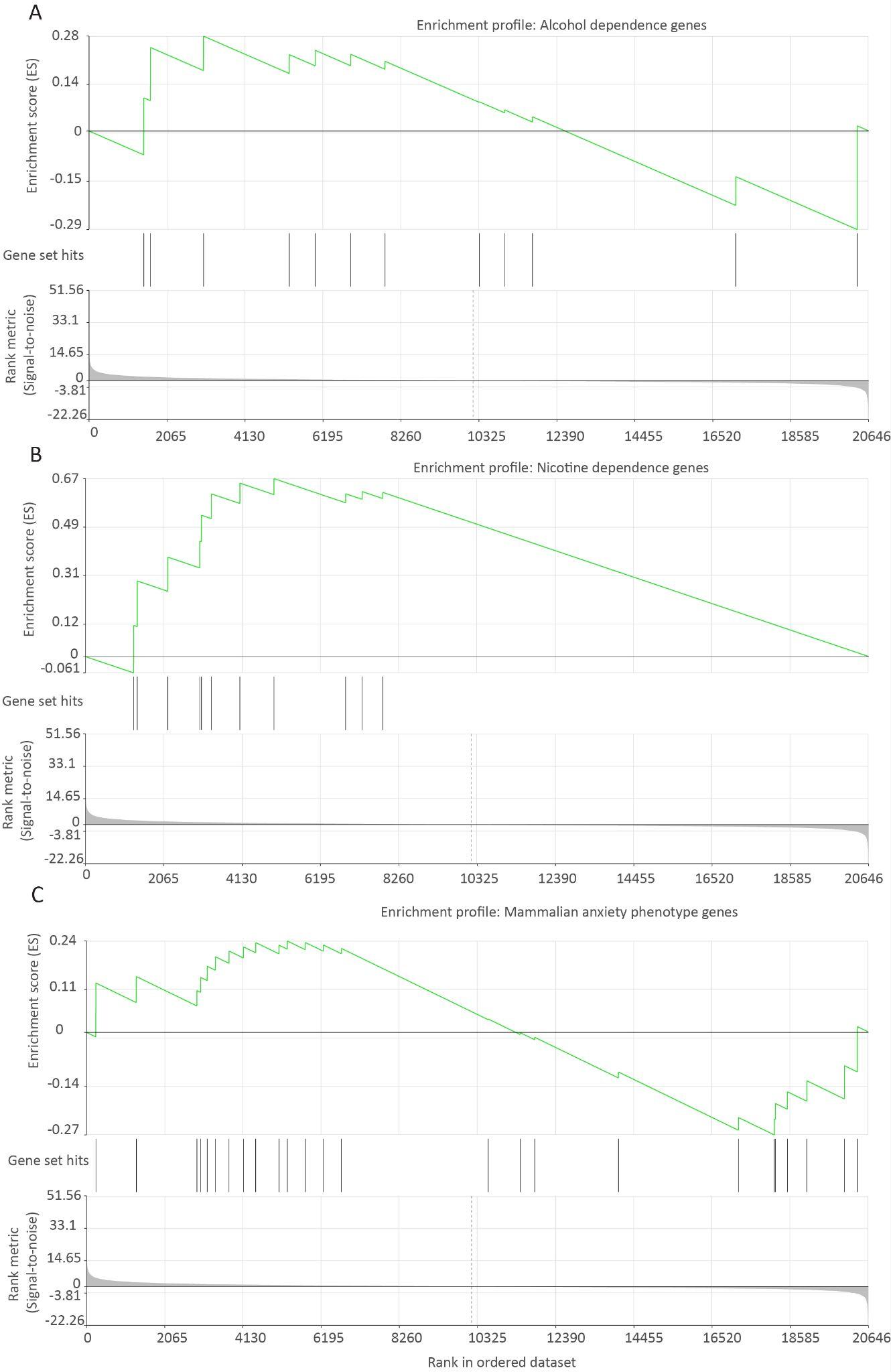

Figure S10. GSEA enrichment profiles for the custom gene sets for nicotine dependence (**A**) and alcohol dependence (**B**) described in table S14, *chrna3^C246X^* vs WT whole brains. See table S15 for statistical analysis.

Table S15. Statistical outputs for figure S10, *chrna3^C246X^* vs WT whole brains.

| **Gene set** | **Gene set size** | **Enrichment score** | **Normalised enrichment score** | **p-value** | **FDR** | **Leading edge size** |
| --- | --- | --- | --- | --- | --- | --- |
| Nicotine dependence | 11 | 0.67 | 1.23 | 0.28 | 0.47 | 8 |
| Alcohol dependence | 12 | -0.29 | -0.88 | 0.6 | 0.72 | 1 |
| Mammalian anxiety phenotype | 25 | -0.27 | -0.75 | 0.75 | 0.83 | 6 |

Table S16. Probe sequences for FISH-HCR. B2 probes were visualised with Alexa fluorophore 647, B3 probes were visualised with Alexa fluorophore 488. P1 and P2 probes of the same gene are paired according to number 1-8.

| **Probe name (gene)** | **Probe sequence** |
| --- | --- |
| B2_P1 (chrna3)_1 | CCTCGTAAATCCTCATCAAACCAAAGATTGGTCTCCATAATCTGG |
| B2_P1 (chrna3)_2 | CCTCGTAAATCCTCATCAAAGGATGAACTCAACACCTCCAAAATC |
| B2_P1 (chrna3)_3 | CCTCGTAAATCCTCATCAAAACCTGGAAATCTCCCACCGCATTGT |
| B2_P1 (chrna3)_4 | CCTCGTAAATCCTCATCAAACTTGAAGATGGCCGGTGGGATCCAG |
| B2_P1 (chrna3)_5 | CCTCGTAAATCCTCATCAAAACGAGCCAAACTTCATGGTGCAGTT |
| B2_P1 (chrna3)_6 | CCTCGTAAATCCTCATCAAATCCCAGAAGTCCTTCAGGTTGATGG |
| B2_P1 (chrna3)_7 | CCTCGTAAATCCTCATCAAACTCCTCACAGCAGTTGTATTTAATG |
| B2_P1 (chrna3)_8 | CCTCGTAAATCCTCATCAAATGATCATGTTGATGGTGTAGAAGAG |
| B2_P2 (chrna3)_1 | CTTATAGTCATTCCAAATGTGTCTCAAATCATCCAGTAAACCGCC |
| B2_P2 (chrna3)_2 | GCTTCCAAATCTTGTTGGACGGCACAAATCATCCAGTAAACCGCC |
| B2_P2 (chrna3)_3 | CGCAGAAGGGCTTTGGTTTTATCGTAAATCATCCAGTAAACCGCC |
| B2_P2 (chrna3)_4 | GTAGGTGACGTCGATCTTGCAGGAGAAATCATCCAGTAAACCGCC |
| B2_P2 (chrna3)_5 | GGTCGATCTTGGCCTTGTCGTAGGTAAATCATCCAGTAAACCGCC |
| B2_P2 (chrna3)_6 | GCGTCGATGATCGTCCATTCTCCGCAAATCATCCAGTAAACCGCC |
| B2_P2 (chrna3)_7 | CAGTGAATACGTGATGTCCGTGTAGAAATCATCCAGTAAACCGCC |
| B2_P2 (chrna3)_8 | TCAGGAAGGAGATGAGAAGACAGGGAAATCATCCAGTAAACCGCC |
| B3_P1(nrp1a)_1 | GTCCCTGCCTCTATATCTTTGTAATCACCCATATGCATTTCTGAG |
| B3_P1(nrp1a)_2 | GTCCCTGCCTCTATATCTTTGCACTCTCGGTCTTCCAGGTCAAAG |
| B3_P1(nrp1a)_3 | GTCCCTGCCTCTATATCTTTTTCCACAATATTTGCCCACCAGCTG |
| B3_P1(nrp1a)_4 | GTCCCTGCCTCTATATCTTTGTCTCGTAGTCGGACACAAACTTGA |
| B3_P1(nrp1a)_5 | GTCCCTGCCTCTATATCTTTGGTGAAGTTCCTGGAACATTCTGGA |
| B3_P1(nrp1a)_6 | GTCCCTGCCTCTATATCTTTTGAACGTGCAGTCCAAATTATTGGG |
| B3_P1(nrp1a)_7 | GTCCCTGCCTCTATATCTTTTGCGTGTCTGGCTCCAGCTCAAAAC |
| B3_P1(nrp1a)_8 | GTCCCTGCCTCTATATCTTTTGGACCAACTCCAGGGAATCCGTCC |
| B3_P2(nrp1a)_1 | AAAATCCTCTGGTTGGGTCCTGGAGTTCCACTCAACTTTAACCCG |
| B3_P2(nrp1a)_2 | GTCTCTCACTTCCACATAGTCATATTTCCACTCAACTTTAACCCG |
| B3_P2(nrp1a)_3 | ACGAGACCACCGGAGATGGAGCGATTTCCACTCAACTTTAACCCG |
| B3_P2(nrp1a)_4 | TAGCGGATGGAGAATCCGGCACCGTTTCCACTCAACTTTAACCCG |
| B3_P2(nrp1a)_5 | GGGCGACTTGATGACTCCGCTGCTGTTCCACTCAACTTTAACCCG |
| B3_P2(nrp1a)_6 | TTTCTGACATCTTAGGAGCAAAGATTTCCACTCAACTTTAACCCG |
| B3_P2(nrp1a)_7 | TATCGGCAGAAGACTCCGGCGGGCGTTCCACTCAACTTTAACCCG |
| B3_P2(nrp1a)_8 | ATTCTGTCCGCAGTATCTGCCGATGTTCCACTCAACTTTAACCCG |
| B1_P1(elavl3)_1 | GAGGAGGGCAGCAAACGGAATCCTCTGCTTCGTTCCGTTTGTCGA |
| B1_P1(elavl3)_2 | GAGGAGGGCAGCAAACGGAAGTTGGCGAACTTTACGGTGATGGGC |
| B1_P1(elavl3)_3 | GAGGAGGGCAGCAAACGGAACAGTGTAGCGGCGAGCGGCTGTCTG |
| B1_P1(elavl3)_4 | GAGGAGGGCAGCAAACGGAAATGGTTATGGGGGAGAATCTGGGGA |
| B1_P1(elavl3)_5 | GAGGAGGGCAGCAAACGGAAGACGAAGATGCACCAGCCGGCTCCA |
| B1_P1(elavl3)_6 | GAGGAGGGCAGCAAACGGAATTGTGACGGCGCCAAAAGGCCCGAA |
| B1_P1(elavl3)_7 | GAGGAGGGCAGCAAACGGAATAGTTGGTCATGGTGACGAAGCCAA |
| B1_P1(elavl3)_8 | GAGGAGGGCAGCAAACGGAACGAGACCTGCAGCACGCGGTCGCCC |
| B1_P2(elavl3)_1 | TTCTGACCGTTCAGGCCCTTGATGGTAGAAGAGTCTTCCTTTACG |
| B1_P2(elavl3)_2 | AGCCTGTCCTGTCTTCTGACTGGGGTAGAAGAGTCTTCCTTTACG |
| B1_P2(elavl3)_3 | AGCGCTGGGTCTGGTGGTGCAGAGGTAGAAGAGTCTTCCTTTACG |
| B1_P2(elavl3)_4 | ACCCCGGCAAGACTAGTCATGCTGTTAGAAGAGTCTTCCTTTACG |
| B1_P2(elavl3)_5 | TTCGTCAGCTTCCGGGGACAGGTTGTAGAAGAGTCTTCCTTTACG |
| B1_P2(elavl3)_6 | TGGTGAAGTCACGGATGACCTTGACTAGAAGAGTCTTCCTTTACG |
| B1_P2(elavl3)_7 | CTGGCGATAGCCATGGCTGCCTCGTTAGAAGAGTCTTCCTTTACG |
| B1_P2(elavl3)_8 | AGCCTTGTGCTGCTTGCTGGTCTTGTAGAAGAGTCTTCCTTTACG |
| B3_P1(chata)_1 | GTCCCTGCCTCTATATCTTTTCTGAACTGGTCCTCTGGGACGAGG |
| B3_P1(chata)_2 | GTCCCTGCCTCTATATCTTTGTAACTTTTTCTGAAGGGTTTCCCC |
| B3_P1(chata)_3 | GTCCCTGCCTCTATATCTTTCTGTTGTTCAAATACATGTCTTCAA |
| B3_P1(chata)_4 | GTCCCTGCCTCTATATCTTTATCACTTTGACCCTTGAAGTTCTGC |
| B3_P1(chata)_5 | GTCCCTGCCTCTATATCTTTGAGCACGTCCATCAATTAGAGCTTT |
| B3_P1(chata)_6 | GTCCCTGCCTCTATATCTTTAAGACCTTGTTGTACTGATCCATAC |
| B3_P1(chata)_7 | GTCCCTGCCTCTATATCTTTTTCAGGCATAACTGTGCTCTTCTGA |
| B3_P1(chata)_8 | GTCCCTGCCTCTATATCTTTGCCGGCGAAAGTTTACCATCACGTC |
| B3_P2(chata)_1 | AAATTTCTCTACAACGGCCTTGGTCTTCCACTCAACTTTAACCCG |
| B3_P2(chata)_2 | AATTGGCCTTTTGTTCACTTCTCTCTTCCACTCAACTTTAACCCG |
| B3_P2(chata)_3 | GGACTGGAGTTGACAGGTAGTGCTATTCCACTCAACTTTAACCCG |
| B3_P2(chata)_4 | AATGAGATTAGCAGCAAATCTGAGGTTCCACTCAACTTTAACCCG |
| B3_P2(chata)_5 | ACTGCCCACGAGCATGCTCTACAGGTTCCACTCAACTTTAACCCG |
| B3_P2(chata)_6 | TTTGTTCCTGGCAGCCTGTAGGACGTTCCACTCAACTTTAACCCG |
| B3_P2(chata)_7 | CTTACATGCCACTATTATATGTTCGTTCCACTCAACTTTAACCCG |
| B3_P2(chata)_8 | GCTGAGTGTAAAGGTCTTTTTCATTTTCCACTCAACTTTAACCCG |

1.	Hu, R., Dai, Y., Jia, P. & Zhao, Z. ANCO-GeneDB: annotations and comprehensive analysis of candidate genes for alcohol, nicotine, cocaine and opioid dependence. *Database* **2018**, (2018).
